# High Throughput Nanopore Sequencing of SARS-CoV-2 Viral Genomes from Patient Samples

**DOI:** 10.1101/2021.02.09.430478

**Authors:** Adrian A. Pater, Michael S. Bosmeny, Mansi Parasrampuria, Seth B. Eddington, Katy N. Ovington, Adam A. White, Christopher L. Barkau, Paige E. Metz, Rourke J. Sylvain, Ramadevi Chilamkurthy, Abadat O. Yinusa, Scott W. Benzinger, Madison M. Hebert, Keith T. Gagnon

**Author notes:** Equally contributing authors (Mansi Parasrampuria, Seth B. Eddington, Katy N. Ovington).

## Abstract

In late 2019, a novel coronavirus began spreading in Wuhan, China, causing a potentially lethal respiratory viral infection. By early 2020, the novel coronavirus, called SARS-CoV-2, had spread globally, causing the COVID-19 pandemic. The infection and mutation rates of SARS-CoV-2 make it amenable to tracking movement and evolution by viral genome sequencing. Efforts to develop effective public health policies, therapeutics, or vaccines to treat or prevent COVID-19 are also expected to benefit from tracking mutations of the SARS-CoV-2 virus. Here we describe a set of comprehensive working protocols, from viral RNA extraction to analysis using online visualization tools, for high throughput sequencing of SARS-CoV-2 viral genomes using a MinION instrument. This set of protocols should serve as a reliable ‘how-to’ reference for generating quality SARS-CoV-2 genome sequences with ARTIC primer sets and next-generation nanopore sequencing technology. In addition, many of the preparation, quality control, and analysis steps will be generally applicable to other sequencing platforms.

## INTRODUCTION

COVID-19, an ongoing global pandemic, has taken the lives of over 400,000 people in the United States (U.S) and approximately 2 million people worldwide in its first year (https://www.worldometers.info/coronavirus/). Effective pandemic response and infection containment have remained challenging (*1*). The ability of viruses to evolve quickly further underscores the need for constant surveillance and monitoring, especially at the genetic level for tracking emergence of novel variants of concern (*2*). Viral genome sequencing can help address these challenges and shed light on the identity and evolution of genetic variants (*3*). Additionally, viral genome sequencing can delineate conserved and mutable regions which would provide valuable insights in the development of effective treatments and vaccines (*4-6*).

Several different approaches exist to sequence SARS-CoV-2 genomes, such as metagenomic sequencing, targeted enrichment sequencing and PCR-amplification based methods (*6*). Metagenomic and target enrichment sequencing are prohibitively expensive to generate large-scale complete genomes and are associated with incomplete coverage and depth (*7*). A solution to address these limitations is a PCR-based amplification which enriches and amplifies the target of interest at the same time (*8*). By utilizing a tiling amplicon scheme with multiplex PCR, it is possible to generate enough coverage and depth to produce complete genome sequences of SARS-CoV-2 in a cost-effective manner (*9*). This approach can even reconstruct complete genomes of samples that have partially degraded viral RNA genomes or low viral load.

Several groups have demonstrated the ability to sequence SARS-CoV-2 using nanopore sequencing with high accuracy at a consensus-level rate and high sensitivity for detecting single nucleotide variants (SNVs) (*10-13*). Our lab adopted and modified protocols initially developed by the ARTIC Network (*9*) to sequence SARS-CoV-2 viral genomes using the MinION nanopore sequencing platform from Oxford Nanopore Technologies (ONT). We present here an entire validated workflow in a 96-well plate format, including a how-to guide for every step from viral RNA extraction to data visualization. These protocols have been validated and benchmarked through the successful sequencing of over 1000 high-quality SARS-CoV-2 genomes from the U.S. state of Illinois (*14*), which are available on the GISAID database and publicly viewable on Nextstrain (https://nextstrain.org/groups/illinois-gagnon-public/ncov/gagnon).

## RESULTS and DISCUSSION

The entire workflow to generate SARS-CoV-2 genomes (**Figure 1**) is performed in a 96-well plate format. All detailed working protocols are provided in the supplemental material and are referenced throughout. The workflow begins with RNA extraction from viral transport media (VTM) containing patient-derived nasopharyngeal (NP) swabs or other potentially other clinical sample sources. Extracted RNA is converted to complementary DNA (cDNA) and viral load approximated by quantitative PCR (qPCR). Following qPCR each sample under a certain cycle threshold (*C*_t_) undergoes two multiplex PCR reactions using ARTIC V3 primer pools that specifically amplify the SARS-CoV-2 viral genome to generate overlapping amplicons products.PCR amplicons can then be optionally visualized by gel electrophoresis. The two separate

**Figure 1.**
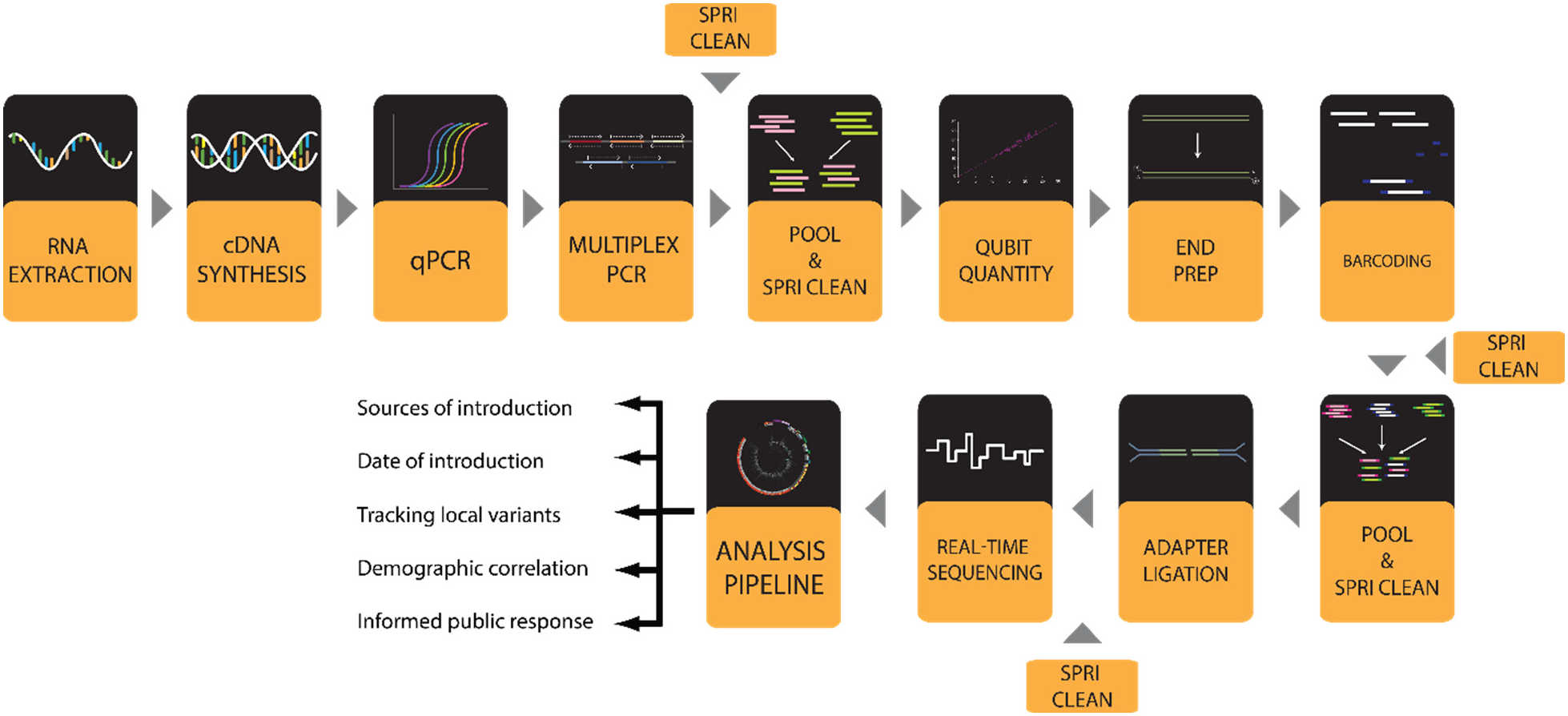
Workflow for SARS-CoV-2 viral genome sequencing with a MinION instrument.

ARTIC PCR product pools for each sample are pooled and the amplicons purified through a clean-up step with paramagnetic Solid Phase Reversible Immobilization (SPRI) beads. The purified PCR amplicons can be optionally quantified by Qubit fluorescence prior to library preparation, which entails an amplicon end-preparation, 96-well sample barcoding, barcoded amplicon pooling and clean-up, and sequencing adapter ligation and clean-up. Sample library is loaded onto a MinION nanopore sequencer (ONT) and sequenced. The bioinformatic analysis to generate whole genome consensus sequences involves basecalling of raw reads, demultiplexing of barcodes, filtering by length, alignment to the SARS-CoV-2 genome, variant calling and consensus sequence generation. Finally, lineage and clade of consensus genomes can be assigned using online tools such as Pangolin and Nextclade (www.pangolin.cog-uk.io) (*15*). Phylogenetic analyses can be performed using Nextstrain (*16*) and results visualized on nextstrain.org. An overview of workflow steps is provided and detailed working protocols for each step are included in supplemental materials.

### Extraction of Viral RNA Genomes

The viral RNA genome is typically extracted from various VTM solutions of clinical COVID-19 samples. Samples should be stored at -80 °C, thawed on ice, and kept cold to maintain viral genome integrity. To increase safety and compliance, our laboratory only receives samples from clinics or public health laboratories that have first been treated with an inactivation solution (*17*) that is compatible with our RNA extraction protocol (see *Working Protocol S1: Viral RNA Extraction*). The RNA extraction is carried out in a BSL-2 class biosafety cabinet. A magnetic bead-based extraction is performed using a 96-well magnetic plate following the manufacturer’s recommended protocol with some minor modifications. These include the use of home-made lysis (inactivation) solution (see Working Protocol S1) in place of, or mixed 1:1 with, the manufacturer’s lysis solution. Additionally, elution is performed in a buffer containing EDTA and an RNase inhibitor to help preserve the quality and integrity of RNA. Once RNA extraction is complete, RNA samples are kept on ice and immediately converted to cDNA. The extracted RNA is then stored in heat-sealed plates at -80°C.

During the RNA extraction it is always recommended to include a negative control containing phosphate buffered saline (PBS) or clean VTM in place of a patient sample in at least one well of each 96-well plate. Negative control samples should procced through the entire workflow and be sequenced if they generate *C*_t_ values in qPCR to determine the extent of contamination. It should be assumed that contaminating amplicons in the negative control might be present in all the samples. Additionally, a positive commercially available sample with a known genetic sequence can be used to validate the workflow at early stages but do not need to be included in every sequencing run. These controls can help identify general cross-contamination issues and ensure efficient extraction. When setting up the workflow for the first time, extracted RNA can be quantified using a spectrophotometer and resolved on denaturing agarose gels or analyzed using a bioanalyzer to evaluate RNA extraction efficiency and RNA integrity. We do not provide protocols for these optional steps.

### Reverse Transcription of Viral RNA to Complementary DNA (cDNA)

Following extraction, RNA is converted to cDNA as soon as possible, preferably without a freeze-thaw cycle, to prevent possible RNA degradation. We have typically used the affordable ABI Hi-Capacity cDNA Synthesis Kit in 96-well format and, in addition to the manufacturer’s recommended protocol, included a small addition of magnesium chloride (MgCl_2_) (see *Working Protocol S2: cDNA Synthesis*). Other reverse transcription kits have not been systematically tested but are expected to be suitable. A constant volume (10 µL) of RNA, half the total cDNA reaction volume, is used as template in all cDNA reactions. Although each sample will have varying concentrations of viral RNA, we have found that this is sufficient to recover genomes from samples with sufficient viral load and does not result in any significant reaction inhibition while reducing hands-on time. Our protocol includes a moderate heating and snap-cooling step in the presence of random primers to improve primer binding (see *Working Protocol S2)*. After synthesis, cDNA can be stored temporarily at 4°C while awaiting qPCR and multiplex ARTIC PCR. For long-term stability, store at -20°C or -80°C after heat-sealing.

### Quantitative PCR (qPCR) to Approximate Viral Load

To quantify approximate viral load in each sample, qPCR is performed using cDNA as template. To reduce costs and maintain consistency, we typically use only one primer-probe set, commercially available from Integrated DNA Technologies (IDT). This primer-probe set, N2, detects the nucleocapsid (N) gene and is identical to one of the primer-probe sets recommended early in the pandemic for qPCR-based patient diagnosis of probable COVID-19 infection (11). We typically use the PrimeTime Gene Expression Master Mix (IDT) and manufacturer’s recommended protocol for qPCR reactions (see *Working Protocol S3: qPCR*). To help standardize quantitation across plates, the same baseline threshold value is set for each run to consistently call *C*_t_ values (**Figure 2A**). In our experience, the *C*_t_ values determined for any given sample are higher than those usually reported (when such data is available) by the facilities that provided patient samples. This is possibly due to a number of factors, including variations in kits used, freeze-thaw cycles, and sample handling. Our recommendation is to avoid freeze-thaw of purified RNA prior to cDNA synthesis to help reduce further loss of viral genome integrity.

**Figure 2.**
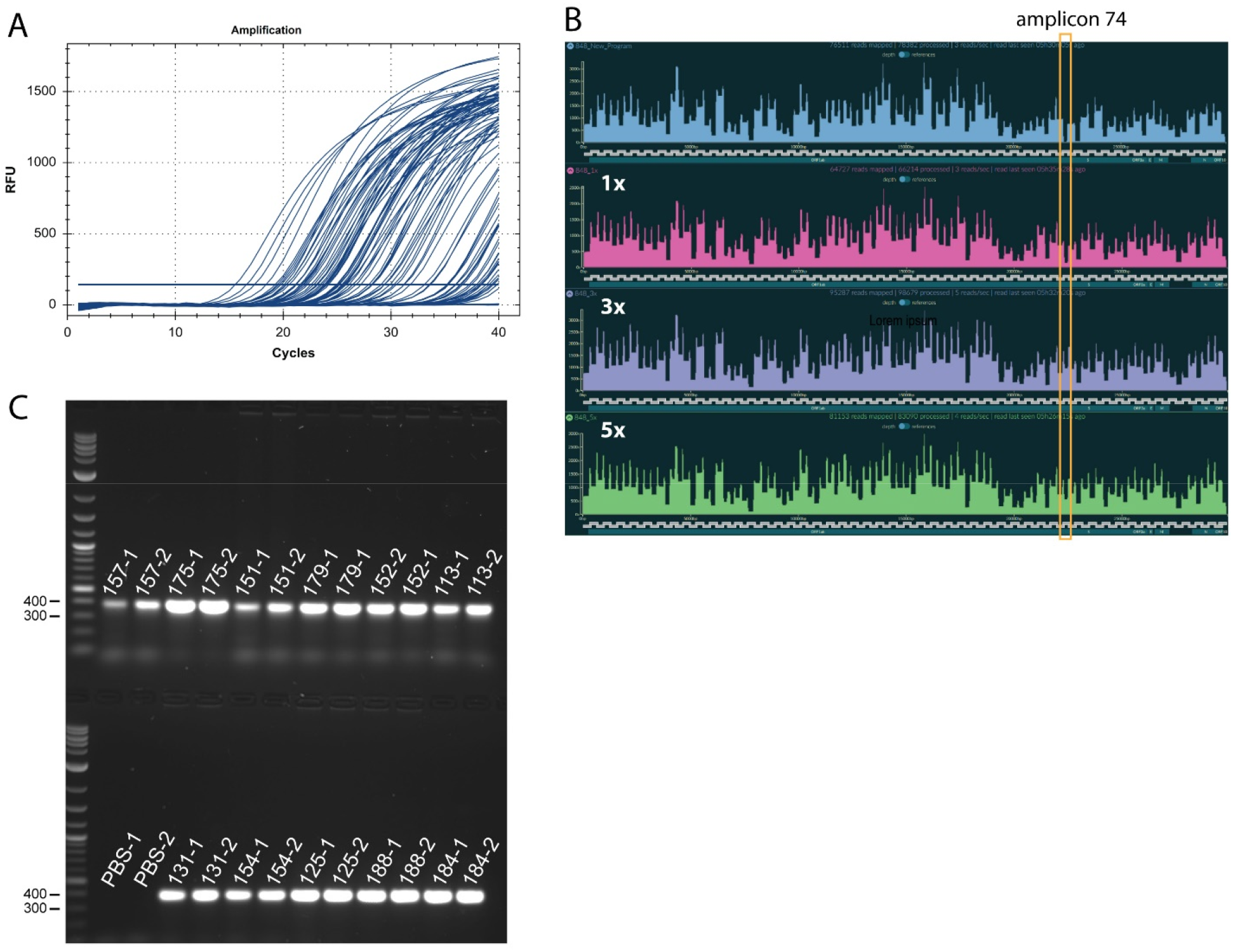
Representative quantitative and ARTIC PCR results. **(A)** Amplification curves from SARS-CoV-2 qPCR assay and IDT 2019-nCoV CDC approved N2 primers and probes. Baseline adjustment was manually set to 200. **(B)** Coverage plot by Rampart with amplicon 74 dropout using ARTIC V3 Primers (top panel/blue plot). Spike-in plots for 1x (15 nM) (second panel/pink plot), 3x (45 nM) (third panel/purple plot) and 5x (75 nM) (bottom panel/green plot) 74_Right and 74_Left are shown for the same sample (*C*_t_ = 20.68). These experiments were conducted at the same time using otherwise identical reagents and protocols. Amplicon 74 region is indicated with an orange box. (**C**) Representative agarose gel electrophoresis of ∼400 bp PCR products amplified using ARTIC nCoV V3 Primers (IDT). Reaction pool 1 (labels with suffixes -1) and pool 2 (labels with suffixes -2) were run side by side with 5 uL of sample on the gel. PBS (no template) extraction controls for pool 1 and 2 show no cross-contamination.

### Multiplex ARTIC PCR to Amplify Viral Genome Sequences

Once qPCR is performed, viral genome cDNA is amplified to increase the copy number of viral genomes. We use two pools of primers designed by the ARTIC Network (*8, 9*), which are commercially available from IDT as the ARTIC V3 primer set. These require two pools of PCR reactions for every sample, with pool 1 having 55 primer sets and pool 2 having 54 primer sets. Two primer pools are used to create overlapping amplicons that reduce interference during PCR and prevent short overlapping products from being preferentially produced. Through this tiling amplicon scheme, complete coverage of the genome is achieved. We also use the Q5 High-Fidelity DNA Polymerase from New England Biolabs (NEB) for PCR reactions (see *Working Protocol S4: Multiplex ARTIC PCR*).

We have optimized the PCR cycling conditions to generate higher and more consistent coverage from higher *C*_t_ samples than can typically be achieved with previously described ARTIC PCR protocols (*8, 9*). Despite optimized cycling conditions, one amplicon is universally known for dropping out: amplicon 74 in primer pool 2. To address this issue, we spiked-in 1-, 3-, or 5-fold more of the primer set for amplicon 74 (which was purchased separately from IDT) and observed improved coverage at 3- and 5-fold molar excess (**Figure 2B**). None of these conditions appeared to reduce coverage of other amplicons from separate samples tested. In fact, coverage of amplicon 76, also often low, seemed to improve as well. Thus, we propose using 3-fold molar excess (45 nM) of amplicon 74 primers over the amount typically present. The details of our optimized ARTIC PCR protocol are found in *Working Protocol S4*.

In principle, quantitation of viral genomes by qPCR should enable precise addition of specific amounts of cDNA for enrichment and amplification in multiplexed ARTIC PCR. However, in practice we have found that this does not improve quality or yield and dramatically increases hands-on time. We have determined that a single defined amount of cDNA volume per ARTIC PCR reaction, up to ¼ of the total reaction volume (5 µL), will provide full genomes up to a *C*_t_ value of 35 in our hands. Thus, we use qPCR to triage samples that are unlikely to yield full genomes. Samples with a *C*_t_ value over 35 using our protocols are usually not processed further. ARTIC PCR reactions can be stored at -20°C or -80°C but are usually kept at 4°C while awaiting clean-up and quantification, described below.

### Pooling and purification of ARTIC PCR Products

Prior to library preparation, pool 1 and pool 2 of ARTIC PCR reactions are combined for every sample and the ∼400 base-pair (bp) amplicons are purified from contaminating PCR reactants using SPRI AMPure XP beads (Beckman-Coulter). When first performing this workflow, it is recommended that 1/5 of the total volume of each pool’s PCR reaction be separately resolved by agarose gel electrophoresis to visualize a specific band for each pool at the correct size of ∼400 bp. (**Figure 2C**). The band intensity can be compared to *C*_t_ values from qPCR to approximate the efficiency of ARTIC PCR and adjust cutoff values for further processing. The remaining pool 1 and pool 2 for every sample are combined and then purified with paramagnetic SPRI beads in a 96-well format following the manufacturer’s recommended protocol (see *Working Protocol S5: SPRI Clean-up of ARTIC PCR*). This protocol requires a 96-well magnetic plate. After this step, DNA can be frozen at -20°C or -80°C but is most often kept at 4°C temporarily until Qubit quantification and end preparation reactions can be performed.

### Fluorescence-Based Qubit Quantification of Pooled and Purified ARTIC PCR Products

The purified PCR pools are quantified using a Qubit 2.0 Fluorometer and Qubit dsDNA High Sensitivity (HS) Assay Kit following the manufacturer’s recommended protocol (see *Working Protocol S6: Qubit Quantification*). This method is specific to double-stranded DNA and is suitable for detection of low levels of DNA. Quantification allows the same amount of the cleaned-up and pooled PCR product for each sample to be used in the next step. This ensures that similar number of reads from each barcode are loaded onto the flow cell and none of the barcoded samples are overrepresented during the sequencing run.. When first performing this workflow, we recommend quantifying all samples with Qubit to quantify the amount of PCR product available and determine the level of recovery with respect to *C*_t_ values from qPCR. However, to reduce time and cost, we routinely only quantitate a representative group of 10-20 samples to ensure expected results are being obtained. We have found that our multiplex ARTIC PCR protocol results in similar concentration for every sample, ∼100 ng/µL, after SPRI clean-up for samples with *C*_t_ values up to 35. If the concentration exceeds the maximum detectable range of Qubit dsDNA HS assay in the previous step, we use an approximate concentration of 100 ng/μL because the concentration of the PCR reaction plateaus at ∼100 ng/μL after 35 cycles.

### Preparation of DNA Ends for Barcoding

Following the approximation of pooled PCR product concentration by Qubit, 60 ng of pooled and SPRI purified PCR amplicons from SPRI clean-up are added to the end preparation reactions designed to create compatible ends of the DNA amplicons for sample barcoding. These are referred to as end prep reactions. The DNA is first end-repaired followed by dA-tailing and inactivation of end-repair enzymes. We use the Ultra II End-Prep kit from NEB following the manufacturer’s recommended protocol with minor modifications (see *Working Protocol S7: Library End Prep*). After this step, DNA can be frozen at -20°C but is most often kept at 4°C for all downstream library preparation steps.

### DNA Library Sample Barcoding and Adapter Ligation

To sequence a full 96-well plate of samples, each sample must be uniquely barcoded and later demultiplexed by allocating reads to samples with the matching barcode. This reduces the overall cost per sample and allows sequencing of up to 96 barcodes on a single run. To achieve this, we use the Native Barcoding Expansion 96 kit (EXP-NBD196) from ONT. We use the NEBNext Ultra II Ligation Master Mix to ligate barcodes to the DNA library (see *Working Protocol S8: Sample Barcoding*). After individual barcoding reactions are performed in a 96-well plate, all samples are pooled and cleaned-up with SPRI beads similar to that described above. A 0.5x ratio by volume of SPRI beads to pooled sample removes free floating barcodes through size selection. This clean-up step is essential to prevent free barcodes from ligating to the incorrect sample during adapter ligation in the next step. This SPRI clean-up requires a microcentrifuge tube magnetic rack. Barcoded samples are stored temporarily at 4°C while awaiting adapter ligation.

After barcoding and clean-up, ONTadapters are then ligated to each amplicon end for the pooled libraries in a single reaction. This reaction sequentially uses Adapter Mix II (AMII) from ONT, NEBNext Quick Ligation Reaction Buffer, and Quick T4 DNA Ligase (see *Working Protocol S9: Adapter Ligation*). The final reaction is then cleaned up using SPRI beads and eluted. Short Fragment Buffer (SFB) is used for the last two clean-up steps rather than ethanol to reduce free barcode carry over and prevent degradation of the adapter-motor protein complex. After elution, the final library is stored at 4°C until Qubit quantification and priming and loading of a nanopore sequencing instrument, such as the MinION from ONT.

### Loading and Running the MinION

For optimal performance on a MinION instrument, 20 ng of adapter ligated library should be loaded onto the flow cell. Overloading the flow cell results in lower throughput. Therefore, a final Qubit quantification is performed similar to that described above. The DNA sequencing library is then prepared using (ONT) reagents following recommended protocols (see *Working Protocol S10: MinION Loading and Running*). The flow cell is then primed using the Flow Cell Priming Kit (ONT) (**Figure 3A**). Priming and loading is described in detail in *Working Protocol S10*. To start the sequencing run, ONT’s MinKNOW software is used which can be downloaded and installed from ONT’s website (https://community.nanoporetech.com/downloads). If real-time sequencing via Rampart is chosen for 96 samples, MinKNOW demultiplexing should be selected. Currently, Rampart only supports real-time sequencing of 24 samples using the 24 Native Barcoding kit (ONT).Once sufficient coverage is met, the sequencing run should be stopped. The flow cell should be washed and stored appropriately following *Working Protocol S11: Flow Cell Wash and Store*.

**Figure 3.**
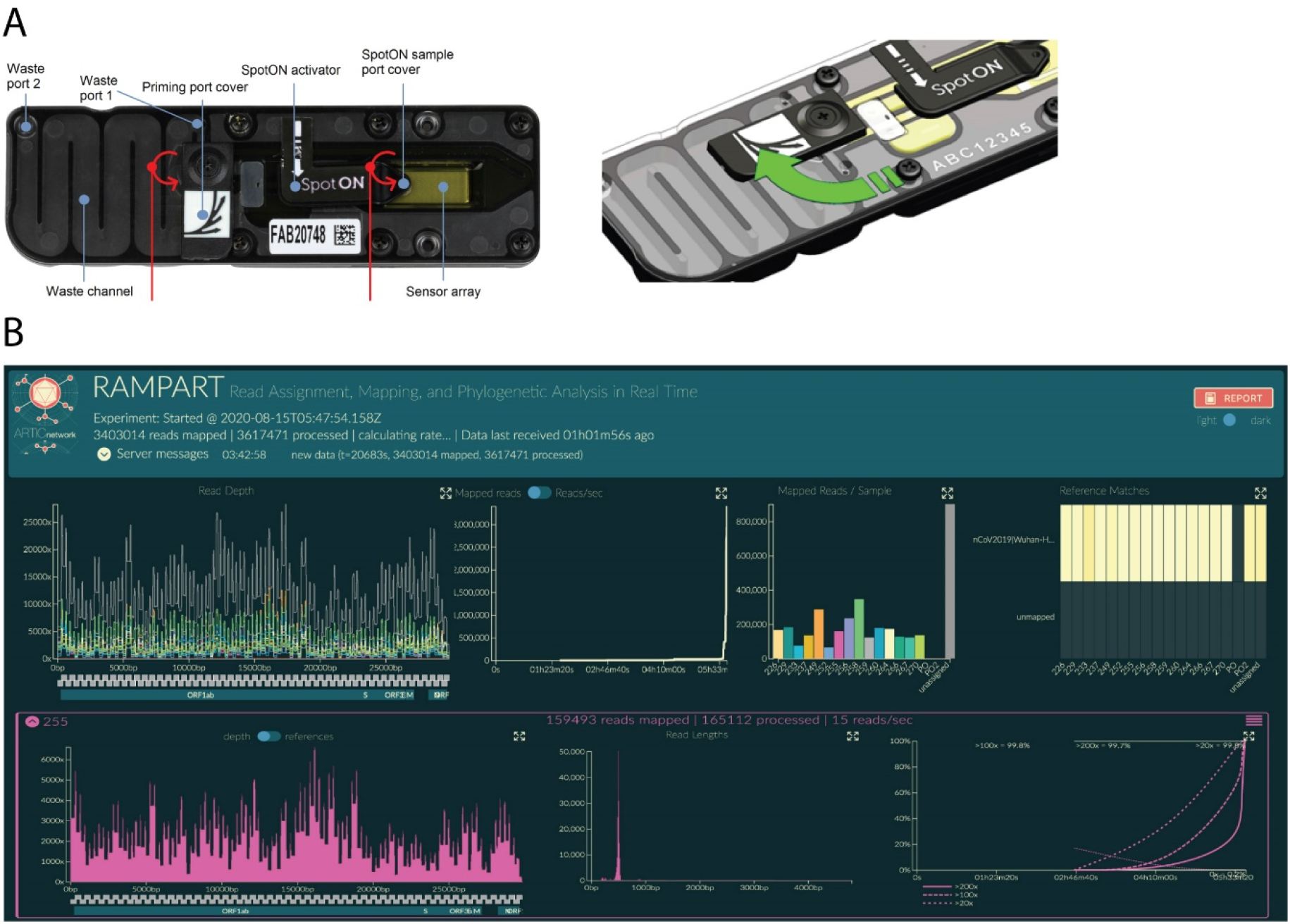
Loading and running the MinION for SARS-CoV-2 genome sequencing. (**A**) Image of the R9.4.1 Flow Cell displaying the different components of the device (left panel). An illustration demonstrating the priming port opening to primer port cover turning clockwise to expose the opening of the priming port (right panel). (**B**) Example of RAMPART display. (Top from left to right): 1) Top left shows coverage across the genomes for all samples. 2) Number of mapped reads through time. 3) Number of reads mapped for each sample. 4) Heatmap showing reads mapped to nCoV2019| Wuhan-Hu-1 (accession MN908947). (Bottom from left to right) 5) The coverage across the genome for an individual sample. 6) Read length distribution for an individual barcoded sample showing the expected peak of ∼400 bp. 7) Percent >20x. >100x and >200x over time.

### Sequencing, Real-Time Visualization, and Data Analysis

Recommendations for desktop requirements and software generally follow the recommendations from ONT. These include a minimal SSD storage of 1 TB, 16 GB of RAM and Intel i7 or Xeon with 4+ cores (https://nanoporetech.com/community/lab-it-requirements). We also recommend using a graphics card from NVIDIA GPU (graphic processing unit) to increase basecalling speed if high accuracy basecalling via Guppy is used.

We recommend implementing Rampart (**Figure 3B**) in order to visualize coverage for each barcoded sample and gather qualitative insight during sequencing runs (https://artic.network/rampart). Rampart can be run concurrently with MinKNOW and be used to determine whether sufficient coverage and depth has been achieved. Since live coverage visualization requires basecalling in near real-time, it is recommended that MinKNOW is set to fast basecalling to quickly generate FASTQ files from fast5 files and gather sequencing insight using Rampart in real time.

A general SARS-CoV-2 targeted amplification bioinformatics pipeline will include basecalling, demultiplexing, trimming, alignment, variant identification and consensus building. In order to reduce false variant calls, a fully validated pipeline should be used (see *Working Protocol S12: Sequencing Data Analysis*). We have implemented the ARTIC bioinformatics pipeline due to its wide use and ONT support (https://artic.network/ncov-2019/ncov2019-bioinformatics-sop.html). Once fast5 files are generated it is highly recommended that high accuracy basecalling is used in order to generate more accurate results of single nucleotide polymorphisms (SNPs). High accuracy basecalling results in better single molecule accuracy but is considerably slower. The basecalled data is then demultiplexed using Guppy with strict parameters to ensure that both barcodes are at both ends of the read. This increases the number of reads binned into the “unclassified” reads but reduces barcode mismatches due to the presence of in silico chimera reads. To remove additional chimeras length filtering is performed using ARTIC guppyplex and reads between 400-700 bp are retained and inputed into the ARTIC bioinformatics pipeline. There are two workflows developed by ARTIC network to generate consensus sequences and identify variants. Nanopolish uses fast5 signal data while the other, Medaka, does not. In our experience both workflows tend to give consistent results with high variant and single nucleotide variant (SNVs) detection accuracy (*10*). We have preferentially used Medaka due to its speed and GPU compatibility. By default, the pipeline minimum coverage is set to 20x to call sites. Otherwise, a masking model applies ambiguous bases (Ns) to sites with lower than the minimum required coverage. After the pipeline has run, a consensus genome sequence is generated which can be concatenated with other consensus genome sequences and further analyzed.

If variants are called from high *C*_t_ samples or partial genomes, the genome sequence should be carefully evaluated and visualized in a graphical viewer. Starting with a very few copies of genomes may lead to artefactual error and call false variants. To reduce artefactual error, it is also recommended to use high fidelity enzymes during cDNA synthesis and PCR. If contamination is observed in the negative control, mutation sites should only be called if sequencing depth greatly exceeds the number of SARS-CoV-2 reads observed in the negative control. To assess amplicon dropouts, we recommend using CoV-GLUE (*18*) which identifies mismatches in sequencing primers/probe sets.

### Phylogenetic Analysis, Variant Calling, and Database Deposition of SARS-CoV-2 Genomes

To process sequenced samples, they must first be properly prepared and formatted (see *Working Protocol S13: Visualizing data in Nextclade and Nextstrain*). A convenient way to observe your sequences and determine the quality is to use the NextClade website (https://clades.nextstrain.org/). Dragging and dropping your sequence FASTA file onto the site will initiate an evaluation process that returns the number of mutations compared to the source SARS-CoV-2 strain, as well as the number of ambiguous nucleotides (Ns) and what ‘clade’ each sequence corresponds to, which are the phylogenetic categories the Nextstrain pipeline separates sequences into. This application also allows searching of the aligned sequences for specific nucleotide or amino-acid mutations (the ‘filter’ button), which is useful for quickly identifying unusual sequences for more careful analysis.

If planning to submit sequences to a global database, such as NCBI or the GISAID Initiative, certain requirements must be met. GISAID requires metadata for each sequence, including collection date and location, the passage history of the sample, and the sequencing technology used. To submit samples in a batch format, more than one at a time, they must be provided with a FASTA file containing all assembled sequences, along with a comma-separated values (.csv) file containing all the metadata. Upon registering and requesting permission for batch upload (https://www.epicov.org/epi3/frontend), a template metadata file is provided explaining the correct formatting. The upload process will do a quality-control pass, alerting users to missing metadata, missing FASTA sequences, or other missing information. Additionally, GISAID is currently checking to ensure that any frameshift mutations present in submitted sequences are correct, and not simply the result of sequencing errors. Submitted sequences with frameshifts must be accompanied with a confirmation email or they will be rejected.

NCBI (https://submit.ncbi.nlm.nih.gov/sarscov2/) has similar requirements. A date and location of collection must be provided. Submitted samples will be processed to trim ambiguous ends, and low-quality sequences (those with too many ambiguous nucleotides, or unacceptably short or long sequences) will be removed.

Finally, if the sequences are going to be placed into a phylogenetic context using the Nextstrain pipeline (https://nextstrain.org/sars-cov-2/), they must also meet requirements there as well. Nextstrain rejects any sample that lacks metadata (ie. time and location of collection, length of sequence) or is too ambiguous. If more than 10% of the sequence is comprised of ambiguous nucleotides, the software will automatically discard it. Nextstrain also estimates the mutation rate of SARS-CoV-2. If samples contain an unlikely number of mutations based on their collection date, these samples will also be excluded from the analysis.

The Nextstrain software can be installed by following their SARS-CoV-2 analysis tutorial (https://nextstrain.github.io/ncov/). Briefly, after the software is installed, all sequence FASTA files must be added in bulk to a sequences.fasta file in the Nextstrain data directory, and the accompanying metadata for those sequences must be added to the metadata.tsv file in the same location. Then a profile must be built that directs the software to focus on those sequences. This can be done using the provided example profiles and is tutorialized in the same link above (see *Working Protocol S13*). Once this is completed, running the analysis is as straight-forward as running a single command. It generates a file which contains a phylogenetic tree of the sequences, a map of collection locations, and a graph of the diversity of mutations in the sampled sequences. Results are in the form of a .json file, created in the auspice subdirectory, which can be viewed by dragging-and-dropping the file onto the Auspice website (https://auspice.us/). More permanent hosting for these analyses can be setup via the Nextstrain website (https://docs.nextstrain.org/en/latest/guides/share/nextstrain-groups.html). An example of our laboratory’s current Nextstrain build with genome sequences from Illinois is shown in **Figure 4**.

**Figure 4.**
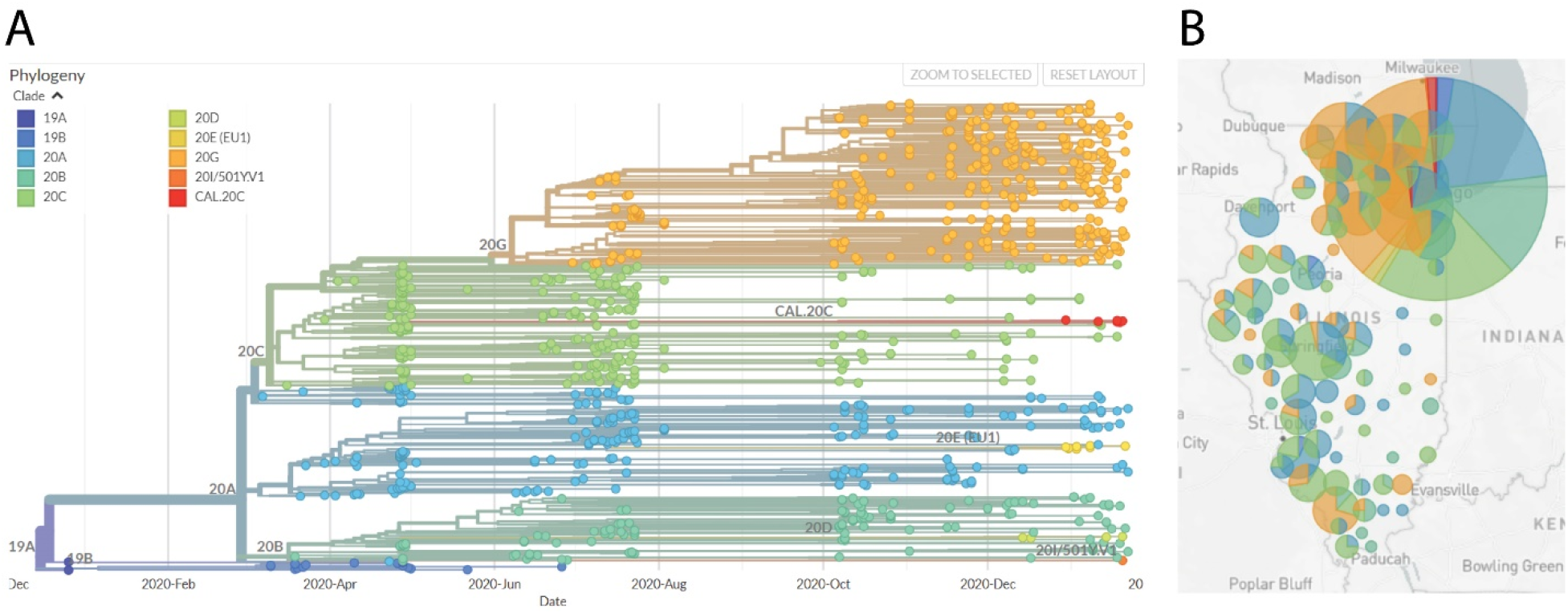
Nextstrain phylogenetic visualization and map view of SARS-CoV-2 genome sequences. **(A)** Phylogenetic tree generated by the Nextstrain pipeline. Sequences are derived from samples taken in Illinois, from April 2020 through January 2021, and sequenced by our laboratory. Clade colors are indicated. (**B**) Map of Illinois showing sample locations, by county. The size of the circle indicates the relative number of sequences derived from that county. The pie chart indicates the proportional distribution of Nextstrain clades at that location. Clade designation colors correlate with those designated in panel A.

## CONCLUSIONS

Here we presented a comprehensive and well-validated set of working protocols for sequencing high-quality SARS-CoV-2 genomes in high throughput from patient samples. These protocols implement the popular ARTIC PCR and MinION nanopore sequencing with 96 samples at a time. We have optimized several steps for efficiency, amplicon coverage and higher recovery, including multiplex ARTIC PCR. Many of the considerations we have covered in these protocols will translate to other sequencing platforms. This workflow and the accompanying supplemental protocols provide a reliable starting point and a comprehensive reference for those seeking to generate SARS-CoV-2 genome sequences using nanopore sequencing technology, especially from the cost-effective MinION instrument.

## Supporting information

Supplemental Working Protocols

## ACKNOWLEDGEMENTS

We thank the Illinois Department of Public Health for access to patient samples. This work was funded by discretionary funds from the Southern Illinois University (SIU) School of Medicine Dean’s Office and the Office of the Vice Chancellor for Research at SIU Carbondale. This work was also funded by the Walder Foundation through a collaborative agreement with the Open Commons Consortium as part of the Chicago CAN initiative.

