## Supplemental Working Protocols for "High Throughput Nanopore Sequencing of SARS-CoV-2 Viral Genomes from Patient Samples"

[†]Equally contributing authors.

### Working Protocol S1: Viral RNA Extraction.

#### Introduction

The first step in the sequencing of SARS CoV-2 virus genomes is by extracting viral RNA from positive SARS-CoV-2 samples. RNA is extracted from inactivated samples in lysis solution using magnetic beads.

#### Materials & Reagents

**Note:** All reagents & materials were estimated for 96 samples. Omega Mag-Bind® Viral RNA Xpress Kit is used for RNA extraction

##### Reagents

- Samples
- TNA Lysis Buffer (25.34 mL)
- Viral Transport Media (VTM, 19.2 mL)
- Mag-Bind Bead Solution (636 µL)
- 100 % Isopropanol (36.464 mL)
- Freshly Prepared 80 % Ethanol (38.4 mL)
- RNA Resuspension Buffer (7.2 µL)

##### Materials

- 96- Well Deep-Well PCR Plate
- Biorad Microseal B
- RNase Decontamination Solution
- 70% Ethanol

##### Equipment

- BSL-2 Hood
- Full PPE
- Thermomixer
- Alpaqua 96-well magnetic platform
- Plate Sealer

#### Procedure

**Caution:** This is the protocol when risk of infection is highest. All steps to be performed in the BSL-2 biosafety cabinet. Full PPE must be worn during the entire extraction process. Samples should only be handled by authorized personnel.

Note: All solutions are made for 106 samples (105%) to ensure sufficient despite pipetting error.

##### *Preparing Working Stock Lysis Buffer*

1. If samples are not already inactivated by Lysis Buffer addition create a working stock to supply 240 µL/sample.
2. Remove Carrier RNA from the -20 °C freezer and allow to thaw out.
3. This is composed by mixing 239 µL TNA Lysis Buffer and 1 µL carrier RNA.
    a. 10 mg/mL tRNA can be substituted in place of carrier RNA.
4. Combine reagents and store in a 50 mL conical tube. Keep at room temperature.

#### *Inactivating Samples in VTM solutions*

1. Aliquot 200 µL of sample in VTM into each well of a 96-well deep-well plate.
2. Add 240 µL of Working Stock Lysis Buffer to each well with sample.
3. Seal the plate with a BioRad Microseal B clear plate sealer.
4. Place the sealed plate in on a thermomixer and tape it down.
    a. Allow to mix at room temperature for 5 minutes at 1100 RPM.

#### *Binding RNA to Mag-Beads*

1. Remove the Mag-Bind bead solution from the 4 °C fridge and thoroughly vortex.
2. Each sample requires 350 µL of the Mag-Bind solution, composed of 344 µL of 100 % isopropanol and 6 µL Mag-Bind beads.
    a. For 96 samples, 36.464 mL and 636 µL of 100 % isopropanol and Mag Bind beads, respectively. Mixed and stored in a 50 mL conical tube.
    b. Mix solution by inversion and shaking of the tube.
3. Remove the seal but do not discard as it will be reused.
4. Aliquot 350 µL of the solution into each well of the plate using a single-channel p1000 pipette.
    a. After the Mage-Bead solution has been added to 12 wells, a full row, cap the 50 mL conical and invert it a few times to mix.

Note: Failing to mix causes the beads to settle out and unequally distribute, impacting purification yields.

5. Reseal the plate with the saved Bio-Rad seal. Place it into the thermomixer, tape down the top, and shake at RT for 10 mins at 1100 RPM.

#### *Separating and Washing Beads*

1. After Removing the plate from the thermomixer, place the plate in the 96-well magnetic platform. Tape it down and all the beads to be pulled to the bottom for 5 - 10 minutes.
2. After this time, if the solution is clear, use a multichannel p200 to slowly and discard the supernatant.
    a. This will take 3 - 4 aspirations and should be pulled from the center of the well, only touching the bottom during the last aspiration.
    b. If a significant amount of beads are aspirated, add the solution back to the well and allow the beads to pellet once more before attempting again.
3. There are three rounds of washing the beads. During the first, remove the plate from the magnetic platform and add 400 µL of RMP Buffer placed in a trough to each sample via a multichannel p200 pipette.
    a. Pipette sample up and down multiple times to ensure beads are well mixed with the RMP Buffer.
4. Seal the plate with the same seal and place into the thermomixer. Tape down the top and mix at room temperature for 10 minutes and 1100 RPM.
5. From the thermomixer, place the plate on the 96-well magnetic platform. Allow the beads to settle at the bottom. Once the solution is clear, using a multichannel p200 set to 200, discard all of the wash buffer.
6. Remove the 96-well plate from the magnetic platform and add 400 µL of freshly prepared 80% ethanol from a trough with a multichannel p200 to each sample.

    a. There is no need to pipette the samples up and down again.
7. Seal the plate with the same seal and place into the thermomixer. Tape down the top and mix at room temperature for 8 minutes and 1100 RPM.
8. Once again, after placing the plate onto the 96-well magnetic plate and letting the solution become clear, slowly aspirate the wash buffer without disrupting the beads.
9. Repeat steps 5 - 9, the second wash, once again.

#### *Elution of RNA from Mag-Beads*

1. Following the final wash remove the plate from the 96-well magnetic platform and allow the beads to dry completely. This can take up to 30 minutes.
2. Add 75 µL of RNA Resuspension buffer (20 mM Tris, pH 7.2, 1 mM EDTA, 0.13 fresh U SUPERase-In) held in a trough to the beads with a p200 multichannel pipette.
    a. Mix by pipetting up and down multiple times after adding to break up clumps.
3. Seal the plate and place into the thermomixer. After taping down the top of it, allow the plate to mix for 20 minutes at 1200 RPM.
4. After the plate has been removed from the 96-well magnetic platform and allow the beads to pull at the bottom.
5. After ~ 5 minutes, once the solution is clear, use a multichannel p200 pipette to move the elution to a new 0.3 mL qPCR plate.
    a. It will most likely be unavoidable to not transfer some beads but they will be removed in the next step.
6. Remove the 96-well deep-well plate from the magnetic platform and tape the 0.3 mL plate with eluted samples to the magnetic platform.
7. After the beads are pulled down, about 3-5 minutes, carefully remove the elution from the wells with a multichannel p200 pipette. Move these to a new 0.3 mL PCR plate. This is the final elution.
8. Cover the plate with a fresh seal, label with experiment number, date, and indicate it is an RNA sample.
9. Store at 4 °C until ready for if cDNA prep is made the same day. For long term storage heat seal the plate and store at -80 °C.

Notify the following team members that you have completed this protocol:

- Make cDNA

### **Working Protocol S2**: cDNA Synthesis.

#### Introduction

The viral genome (RNA) is reverse transcribed to complementary DNA (cDNA) using random primers. The resulting cDNA can then be used for qPCR and ARTIC PCR.

#### Materials & Reagents

**Note:** All reagents & materials are estimated for 96 samples. The cDNA synthesis kit used is ABI High Capacity cDNA Reverse Transcription Kit (Catalog #: 4368813).

##### Reagents

- 10x RT Buffer (212 µL)
- 10x RT Random Primers (212 µL)
- 25x dNTP Mix, 100 mM (85 µL)
- MultiScribe RT, 50 U/µL (105.6 µL)
- $MgCl_2$, 50 mM (21.12 µL)

##### Materials

- 96-well PCR plate (1)
- Multichannel pipettes
- 50 mL flat bottom tube (1)
- Electronic Pipettor
- Plate Seals (1)
- Trough (Integra)
- 96-well Aluminium Block

##### Equipment

- 96-Well Heat Block
- Plate Sealer
- Thermal Cycler

**Note:** cDNA synthesis should be done on a clean bench to avoid any chance of contamination. Therefore, decontaminate your bench with RNAse decontamination solution and then 70% ethanol. Use filtered tips and sterile labware to reduce the chance of contamination.

#### Procedure

##### *Plate Setup*

Obtain the 96-well plate of the RNA. Convert it to a layout that will represent this cDNA plate.

##### *Prepare Master Mixes*

**Note:** Enough master mix is made for 106 samples to account for pipetting error.

1. Thaw 10X RT Buffer, 10X RT Random Primers, 25X dNTP mix (100 mM) and $MgCl_2$ (50 mM) at room temperature. Mix by vortexing, centrifuge down using a benchtop centrifuge and place on ice.
2. Gently flick the Multiscript RT (50 U/µL) several times, spin down and immediately place on ice.
3. Set up the 96-well heat block to 65 °C prior hand to heat up.

### cDNA Synthesis

4. Prepare the two master mixes in separate 1.5 mL microcentrifuge tubes by combining the reagents in the following order.
    a. After adding each reagent to the reaction, mix by pipetting up and down 3 times. Be sure to switch tips between each sequential addition.

| Master Mix # 1 Reagents | Volume |
|---|---|
| **Nuclease-Free Water** | 212 µL |
| **10x Random Primers** | 212 µL |
| **25x dNTPs Mix, 100 mM** | 85 µL |
| **Master Mix # 1 Total** | **509 µL** |

| Master Mix # 2 Reagents | Volume |
|---|---|
| **10x RT Buffer** | 212 µL |
| **Multiscript RT (50 U/µL)** | 105.6 µL |
| **MgCl2 (50 mM)** | 21.12 µL |
| **Master Mix # 2 Total** | **338.72 µL** |

5. Retrieve the RNA plate from the -80 °C freezer and immediately place on ice. Thaw the RNA on ice and keep it sealed until it is ready to be added to the reaction plate.
6. Vortex and spin down Master Mix # 1, then add 6.8 µL of it into each well of a 0.3 uL 96-Well PCR plate on ice.
    a. Adding of the master mix can be done by multichannel pipette or single channel pipette.
7. Using a multichannel pipette transfer 10 µL of RNA into their respective wells containing Master Mix # 1 from previous step.
8. Pipette up and down to mix the reaction.
9. Place the PCR plate on the heating block and allow it to incubate for five minutes.
10. Immediately following the incubation period, place the PCR plate by placing it into the 96-well aluminum block on ice for at least one minute.
11. Add 3.2 µL of Master Mix # 2 to each well on ice and pipette up and down to mix and spin the plate.
12. Use adhesive or heat seals and make sure that each well is properly sealed to prevent evaporation.

#### *Thermocycling Plate*

1. Place the PCR plate in the thermal cycler and run at 25 °C for 10 minutes, 37 °C for 2 hours, 85 °C for 5 minutes and hold at 4 °C.
2. Label the RNA plate, heat seal it and store at -80 °C.

Notify the following team members that you have completed this protocol:

- qPCR

### Working Protocol S3: qPCR.

#### Introduction

qPCR is used in order to confirm positive samples and determine which samples can result in full or nearly full genomes from the workflow. Any sample with a cycle threshold ($C_t$) value exceeding a threshold $C_t$ will not proceed through the workflow. This reduces cost and hands-on time. In addition to chances of genome retrieval, the $C_t$ value provides insight on the viral load.

#### Materials & Reagents

**Note:** All reagents & materials are estimated for 96 samples, scale as needed.

##### Reagents

- Nuclease-Free water (318 µL)
- 2X IDT PrimeTime (1060 µL)
- 10 µM N2 Forward Primer (106 µL)
- 10 µM N2 Reverse Primer (106 µL)
- 2.5 µM N2 Probe (106 µL)

##### Materials

- Bio-Rad 96 Well PCR Plate
- 96-Well Plate Seals

##### Equipment

- Plate Sealer
- qPCR Machine

#### Procedure

##### *Plate Setup*

Obtain the 96-well plate of the cDNA layout from the Z-drive. Convert it to a layout that will represent this qPCR plate.

##### *Master Mix Assembly*

**Note:** qPCR should be done on a clean bench to avoid any chance of contamination. Therefore, decontaminate your bench with RNAse decontamination solution and then 70% ethanol.

1. Thaw 2X PrimeTime, 10 µM Forward Primer, 10 µM Reverse Primer, and 2.5 µM Probe Briefly vortex, and spin down briefly before placing on ice.

**Note:** Avoid prolonged exposure of reference dye to light.

2. Prepare the master mix in a 2 mL microcentrifuge tube tube by combining the reagents in the following order, stopping after the 2.5 µM Probe.
    a. After adding each reagent to the reaction, mix by pipetting up and down 3 times. Be sure to switch tips between each sequential addition.

**Note:** The master mix is for 96 samples, adjust as needed.

| Reagents | Volume |
|---|---|
| **Nuclease-Free Water** | 318 µL |
| **2X PrimeTime** | 1060 µL |

| | |
|---|---|
| **10 µM Forward Primer** | 106 µL |
| **10 µM Reverse Primer** | 106 µL |
| **2.5 µM Probe** | 106 µL |
| **Sub Total** | 1696 µL |
| **Sample cDNA** | 4 µL / sample |
| **Final Volume** | **20 µL** |

3. Retrieve the cDNA plate from the 4 °C fridge. Allow to thaw, vortex and spin briefly. Place the plate directly on ice.
4. From the cDNA plate, add 4 µL of the cDNA to a 96-Well qPCR Plate, corresponding with their position on the plate layout.
5. Add 16 µL of the master mix to the well containing cDNA.
   a. After adding each reagent to the reaction, mix by pipetting up and down 3 times. Be sure to switch tips between sample addition.
6. Seal the PCR plate, making sure that each well is well sealed to prevent evaporation. Vortex the plate briefly and spin down.
7. Reseal the cDNA plate, label and place in 4 °C or -20 °C for longer term storage.

##### *Thermocycling Plate*

Run the following thermocycler conditions:

| **Step** | **Temp** | **Time** |
|---|---|---|
| 1- Polymerase activation | 95 ℃ | 3 min |
| 2- Denaturation | 95 ℃ | 10 sec |
| 3- Annealing | 55 ℃ | 30 sec |
| 4- Plate Read | Read plate ***FAM** | |
| Repeat steps 2-4 for 39 cycles. | | |

a. This program will run for ~ 1 hour.

##### *Data Retrieval*

1. Set the baseline threshold at 200 and export the Ct values as xls or csv file.

### Working Protocol S4: Multiplex ARTIC PCR.

#### Introduction

In order to have enough material to obtain sufficient reads during sequencing, cDNA is exponentially amplified using PCR. This multiplex amplification protocol utilizes the V3 pool of ARTIC Primers which contains 218 unique primers, in two alternate pools, to produce ~ 400 base-paired amplicons.

#### Materials & Reagents

**Note:** All reagents & materials are estimated for 48 samples, scale as needed. Primer scheme: https://github.com/artic-network/artic-ncov2019/tree/master/primer_schemes/nCoV-2019/V3

##### Reagents

- 5X Q5 Reaction Buffer (480 μL)
- Q5 High-Fidelity Polymerase (24 μL)
- ARTIC Primer Pool 1 Mix (IDT) (384 μL)
- ARTIC Primer Pool 2 Mix (IDT) (384 μL)
- 2.5 mM dNTP Mix (192 μL)
- 74_Left and 74_Right Primers (IDT)

##### Materials

- 96-Well PCR Plate
- 96-Well PCR

##### Equipment

- Plate Sealer
- Thermal cycler

#### Procedure

##### *Plate Setup*

Obtain the 96-well plate layout containing qPCR results. Create a new layout that removes any sample with a $C_t$ value greater than the choosen threshold. This will represent the ARTIC PCR plate. If more than 48 samples are being run, pool 1 and 2 will require separate plates.

##### *Master Mix Assembly*

**Note:** PCR is extremely susceptible to contamination, work on a clean bench and decontaminate your bench with RNAse decontamination solution and then 70% ethanol. Appropriate precautions should be taken to reduce the chances of contamination.

1. Thaw 5X Reaction Buffer, Primer Pools 1 & 2 and 2.5 mM dNTP mix. Mix at room temperature by vortexing, pulse centrifuge using a benchtop centrifuge and place on ice.
2. Gently flick the Q5 High-Fidelity Polymerase multiple times and immediately place on ice.
3. Dilute 100 Micromolar (μM) primer pools 1:10 in nuclease free water, to generate 10 μM primer stocks.
    a. Make several 10 μM stocks in case of contamination or degradation.
4. Spike in 3x 15 nM of the 74_Right and 74_Left primer into 10 μM primer pool 2.

**Note:** The ARTIC V3 primers are used at a final concentration of 15 Nanomolar (nM) per primer. In this case V3 pools have 110 primers in pool 1 and 108 primers in pool 2. Our initial sequencing analyses constantly observed a drop out in amplicon 74. To overcome this, we spike in primer pair 74 at 3x in the 10 µM primer pool 2.

5. Prepare the pool 1 and pool 2 master mixes in a pre-PCR hood. Separate microcentrifuge tubes by combining the reagents in the following order
    a. After adding each reagent to the reaction, mix by pipetting up and down 3 times. Be sure to switch tips between each sequential addition.

**Note:** The master mix is for 48 samples for each pool, adjust as needed.
**Note:** Enough master mix is made for 52 samples to account for pipetting error.

| Reagents | Pool 1 Volume | Pool 2 Volume |
|---|---|---|
| **Nuclease-Free Water** | 455 µL | 455 µL |
| **5X Q5 Reaction Buffer** | 260 µL | 260 µL |
| **Pool 1 Primer Mix** | 208 µL | 0 µL |
| **Pool 2 Primer Mix** | 0 µL | 208 µL |
| **2.5 mM dNTP Mix** | 104 µL | 104 µL |
| **Q5 High-Fidelity Polymerase** | 13 µL | 13 µL |
| **Sub Total** | 1040 µL | 1040 µL |
| **Sample cDNA** | 5 µL / sample | 5 µL / sample |
| **Total** | **25 µL** | **25 µL** |

6. After making the master mixes put all reagents away and take out the cDNA plate to thaw.
7. Vortex the plate and spin down. Place the plate on ice.
8. From the cDNA plate, add 5 µL of sample cDNA to the pool 1 and pool 2 into the reaction PCR plate, corresponding with their position on the plate layout.
9. Reseal the cDNA plate, label and store at -20 °C.
10. Add 20 µL of Pool 1 Master Mix and Pool 2 Master Mix to the wells containing samples in plate 1 and plate 2, respectively.
    a. After adding each reagent to the reaction, mix by pipetting up and down 3 times. Be sure to switch tips between addition.
11. Seal the PCR plate, making sure that each well is tightly sealed to prevent evaporation. Vortex gently and spin down.

##### *Thermocycle Program*

| Step | Cycle Step | Temp | Time | Number of Cycles |
|---|---|---|---|---|
| 1 | Heat activation | 98 °C | 30 sec | 1 |
| 2 | Denaturation | 94 °C | 16 sec | 20 |
| 3 | Annealing | 65-63 °C Touchdown (-0.1 °C each cycle) | 5 min | 20 |
| 4 | Denaturation | 94 °C | 16 sec | 15 |
| 5 | Annealing | 63 °C | 5 min | 15 |
| 6 | Hold | 4 °C | ∞ | |

    a. This program will run for ~ 3.5 hours.
2. Label, date and place the plates at a 4 °C.

Notify the following team members that you have completed this protocol:

- SPRI Pool and Clean up

### Working Protocol S5: SPRI Clean-up of ARTIC PCR.

#### Introduction

The clean-up is performed by using AMPure XP SPRI beads to remove leftover PCR contaminants (dNTPs, salts, primers, etc.) from the two previously performed PCR reactions. Leftover contaminants may lead to poor DNA library end-prep and barcoding efficiency. The advantage of using SPRI beads is their ability to size select DNA depending on the ratio of bead to sample volume used. Expected recovery is 60-80%.

#### Materials & Reagents

**Note:** All reagents & materials were estimated for 96 samples.

##### Reagents

- AMPure XP beads (4 mL)
- 100% Ethanol (40 mL)
- Omega Elution Buffer (EB) (2.9 mL)
- ddH2O

##### Materials

- 96-well PCR plate (2)
- 96-well magnetic separator
- Multichannel pipettes
- 50 mL flat bottom tube (1)
- Electronic Pipettor
- Plate Seals (2)
- Razorblade (2)
- Trough

#### Preparation

**Note:** Amplicon clean up should be done on a clean bench to avoid any chance of contamination. Therefore, decontaminate your bench with RNAse decontamination solution and then 70% ethanol.

**Note:** You should have a dedicated labeled trough for each reagent. ddH2O, 70% ethanol between each use and dry completely. The trough may be placed at 37 °C to dry faster.

##### *AMPure XP Beads*

You can prepare the beads up to 3 days in advance and scale up accordingly.

1. Resuspend the stock AMPure XP Beads by vortexing for 20 seconds. The solution should be a homogenous brown color.
2. Using an electronic pipettor, remove 4 mL of AMPure XP beads and place it in a clean trough on ice. As soon as you are done, place the stock beads directly back into the 4 °C refrigerator.
3. Using an 8-channel P200 multichannel pipette set to 200 µL, mix the beads in the trough by pipetting up and down 5 times.
4. Immediately, set the multichannel pipette to 40 µL and transfer the beads from the trough to the 96-well PCR plate.
5. Between each transfer, mix the beads by pipetting up and down 2 times.

    a. The beads can quickly settle at the bottom of the trough.
6. After transferring the beads into all 96 wells, seal, label and store the plate at 4 °C or use immediately. This plate will be referred to as the 96-well clean-up plate from this point on.
7. Remove any residual AMPure XP beads from the trough into a labeled and dated 1.5 mL microcentrifuge tube and place at 4 °C for use next time beads are prepared.

##### *80% Ethanol*

Prepare 80% ethanol fresh before use.

Recipe for 80% EtOH
In a 50 mL flat bottom tube:
1. Label the tube.
2. Pour 40 mL 100% ethanol.
3. Add 10 mL of ddH2O.
4. Mix by inversion.

#### Procedure

You will begin this procedure with two 96-well plates containing the PCR product from the previously performed PCR reaction. The plates will be referred to as PCR pool 1 plate and PCR pool 2 plate.

##### *Pooling of PCR Products*

**CAUTION:** Be very careful when pooling PCR products. Avoid cross-contaminating neighboring wells with small droplets.

**Note:** If each pool should is to be resolved on an agarose gel individually, do NOT pool into PCR pool 2 plate. Pool 20 µL of PCR pool 1 and 20 µL PCR pool 2 into the respective wells of the 96-well clean up plate.

1. Remove a prepped 96-well clean-up plate containing beads from 4 °C or prepare a 96-well clean-up plate by adding AMPure XP beads as described in "***AMPure XP Beads***," and place it on a cleaned benchtop.
2. Spin down each PCR product plate of pool 1 and pool 2 at 2,000 RPM for 30 seconds.
3. Place both PCR pool plates (pool 1 and pool 2) into plate holders.
4. Cut the seal between each row of both PCR pool plates using a razorblade so that it can be removed one row at a time.
5. Use an 8-channel P200 pipette and set it to 25 µL (the reaction volume is 25 µL).
6. Carefully, remove the seal to expose row 1 of PCR pool 1 plate and PCR pool 2 plate.
7. Transfer 25 µL from row 1 of PCR pool 1 plate to the corresponding wells of row 1 of the PCR pool 2 plate (Fig 1) and mix by pipetting up and down 3 times.

   **Note:** Be certain that all the liquid from PCR pool 1 plate has been transferred into PCR pool 2 plate.

8. Set the 8-channel P200 pipette to 40 µL and transfer 40 µL of the pooled product from PCR pool 2 plate into the corresponding wells of row 1 96-well clean-up plate (Fig 1) and mix by pipetting up and down.

   **Note:** After transferring 40 µL into the 96-well clean-up plate there should be 10 µL of pooled PCR product left for each sample in the PCR pool 2 Plate to run on the gel.

9. Remove the seal from both PCR pool plates to expose the next row.
10. Repeat steps 5-9, until all of the pooled reactions are transferred from PCR pool 2 plate into the 96-well clean-up plate.
11. Cover the 96-well clean-up plate using a P1000 pipettor tip box lid.
12. Incubate the 96-well clean-up plate at room temperature for 15 minutes.
13. Seal PCR Pool 2 plate containing 10 µL for agarose gel electrophoresis. Label, date and place in 4 °C refrigerator.

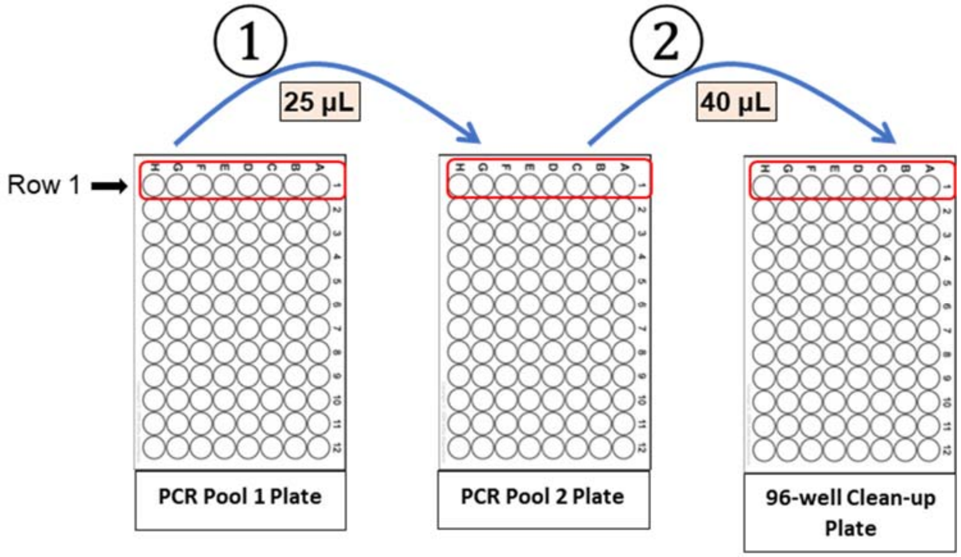


**Figure 1**

##### *Purify the Pooled Reactions*

**CAUTION:** Be very careful not to touch the pelleted beads when removing the supernatant by inserting the tips vertically to prevent the tips from touching the periphery of the wells where beads would be localized.

1. After incubating the 96-well clean-up plate for 15 minutes place it on the magnet for 5 minutes.
2. Check whether the solution became clear. If not, continue to keep the plate on the magnet.
3. Carefully discard the supernatant using a P200 multichannel pipette without disturbing the beads.

**CAUTION:** Be very careful not to touch the pelleted beads when removing the supernatant. Removing any of the beads will drastically decrease yield.

4. While keeping the plate on the magnet, wash the beads with 200 µL of 80% ethanol without disturbing the beads.
5. Keep the plate on the magnet for 3 minutes.
6. Carefully discard the ethanol using a multichannel pipette while the plate stays on the magnet. This ends the first ethanol wash.
7. Repeat steps 4-6 (80% ethanol wash) for a second ethanol wash.
8. Spin down the 96-well clean-up plate at 2,000 RPM for 30 seconds.

9. Place the 96-well clean-up plate on the magnet.
10. Use a P10 single channel pipette to aspirate any residual ethanol you observe.
11. Remove the plate from the magnet and place onto the benchtop.
12. Using a P200 multichannel set to 30 µL, add 30 µL of Omega EB to each of the wells to elute the DNA and mix by pipetting up and down 5 times.
13. Change tips and continue until all of the beads in the 96-well clean-up plate have been resuspended.
14. Cover the plate using a P1000 tip box lid and incubate for 2 minutes at room temperature.
15. Spin down the plate containing the resuspended DNA at 2,000 RPM for 15 seconds.
16. Place the plate on the magnet for 4 minutes.
17. Using a P200 multichannel pipette, carefully remove and retain 30 µL of the eluate containing the DNA library per well into a new 96-well PCR Plate.
    a. Be careful not to remove any beads.
    b. Be careful not to change the plate layout.

**CAUTION:** Do NOT discard the supernatant as it contains the library DNA.

18. Seal, label and date the plate containing 30 µL of the eluate. Place in 4 °C refrigerator.
    a. This plate will be used for quantification and end-prep reaction.

Notify the following team members that you have completed this protocol:

- Run PCR products on agarose gel
- Quantify DNA Samples on Qubit
- Library End Prep

### <u>Working Protocol S6</u>: Qubit Quantification.

#### Introduction

The Qubit dsDNA HS assay uses fluorescent dyes that bind specifically to dsDNA providing a more accurate quantification because it only measures the target of interest (i.e. dsDNA). This assay relies on a 2-point curve by reading the two standards for calibration every time measurements are taken. The assay is designed to quantitate 0.2 – 100 ng in a 200 µL assay.

#### Materials & Reagents

##### Reagents

- Qubit dsDNA HS Buffer
- Qubit dsDNA HS Reagent (Dye)
- Qubit dsDNA HS Standard #1
- Qubit dsDNA HS Standard #2

##### Materials

- Eppendorf Tube
- Qubit assay tubes
- Qubit 2.0 Fluorometer

#### Procedure

**Note:** For best results, store the Qubit dsDNA HS Buffer at room temperature. Store the Qubit standards and Qubit Reagent (Dye) at 4°C. Ensure that all assay reagents are at **room temperature** before you begin.

**Note:** The kit comes with 2 standards (standard 1 and standard 2), reagent (dye) and dsDNA HS Buffer.

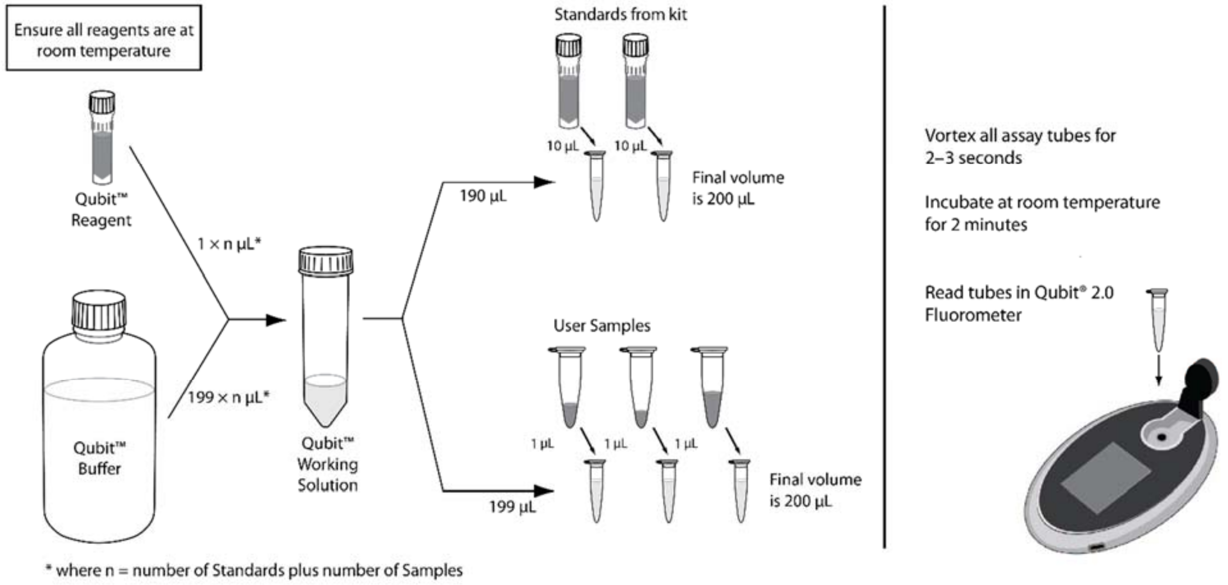

##### *Qubit Reaction Preparation*

1. Remove the Qubit standards and reagent (dye) from 4°C and let them equilibrate to room temperature.
2. Flick the DNA samples to be quantified and briefly centrifuge.
3. Determine the total number of samples + two standards (included in the kit).
4. Set up two Qubit Assay tubes for the two standards and one tube for each sample to be quantified. Label the tube lids.
   a. The Qubit Assay tubes are specific clear thin-wall 0.5 mL tubes designed only to be used with the Qubit fluorometer.
5. Make sufficient Qubit working solution (WS) in a single microcentrifuge tube for the total number of reactions (standards and samples), by combining 1 μL Qubit dsDNA reagent (dye) to 199 μL Qubit dsDNA buffer for each reaction.
6. For example, for 4 samples, prepare enough working solution (WS) for a total of 6 reactions (4 samples + 2 standards = 6 reactions). Add:

| # reactions | 1X | 6X |
|---|---|---|
| **dsDNA Buffer** | 199 | 1,194 |
| **dsDNA Reagent** | 1 | 6 |
| **Total WS (μL)** | 200 | 1,200 |

7. Mix the Qubit working solution (WS) by vortexing for ~4 seconds and briefly centrifuge.
8. Add 190 μL of Qubit working solution (WS) to each of the Qubit assay tubes used for standard 1 and standard 2.
9. Add 10 μL of each Qubit standard to the appropriate tube, mix by vortexing for ~4 seconds and briefly centrifuge.
10. Add 199 μL of Qubit working solution (WS) to each of the Qubit assay tubes used for samples to be quantified.
11. Add 1 μL DNA sample, mix by vortexing for ~4 seconds and briefly centrifuge.
    a. Visually inspect the P10 pipette tip when sample has been aspirated to make sure ~1 μL of sample is dispensed into the Qubit assay tube.
12. Close the lids tightly and allow all tubes to incubate at room temperature for 2 minutes.

##### *Qubit Quantification*

1. On the Home screen of the Qubit Fluorometer, press **DNA**, then select **dsDNA High Sensitivity** as the assay type.
2. The "Read New Standards?" screen is displayed. Press **Yes.**
3. Insert Standard #1 into the Sample Chamber, close the lid and press **Read.**
4. Insert Standard #2, close the lid and press **Read.** Calibration of Qubit is now complete.
5. Choose **Sample** to go to the sample screen.
6. Insert a sample into the sample chamber, close the lid and press **Read.**
7. A result will display on the screen. Press **Calculate Stock Conc.**
8. Using the volume roller wheel, select **1 μL.**
9. Change the units in which the original sample concentration is displayed, press **ng/mL.**
10. Select **ng/μL.** **Record** the Initial Conc.: value for the sample in an excel sheet/notebook.
11. Save the data by pressing **Save.**
12. Repeat for all DNA samples by pressing **Read** once a new sample has been inserted.

**Note:**

- If the sample concentration is too high, dilute the sample 1:1 by volume with Qiagen elution buffer (EB).

### <u>Working Protocol S7</u>: Library End Prep.

#### Introduction

The end-prep reaction is required to create compatible ends of the DNA amplicons for the next step of DNA library preparation. The DNA is first end-repaired at a lower temperature (20 °C) and then, subjected to a higher temperature (65 °C) to promote dA-tailing and inactivate end-repair enzymes. Both end repair and dA-tailing reactions are performed in one tube with an enzyme mixture.

#### Materials & Reagents

**Note:** All reagents & materials were estimated for 96 samples.

##### Reagents

- NEBNext Ultra II End-Prep Enzyme Mix
- NEB Ultra II End-Prep Buffer

##### Materials

- 96-well PCR plate
- Multichannel Pippetes
- Plate Seal
- PCR Tubes
- Thermal cycler
- Trough

##### *End-Prep Master Mix*

**Note:** End prep up should be done on a clean bench to avoid any chance of contamination. Therefore, decontaminate your bench with RNAse decontamination solution and then 70% ethanol.

**Note:** You should have a dedicated labeled trough if making a Master Mix. If reusing trough clean using ddH2O, 70% ethanol and ddH20 again between each use and dry completely. The trough may be placed at 37 °C to dry faster.

**Important:** For optimal efficiency of the end-prep reaction, use ~245 fmol (65 ng for 400 bp amplicons) of the cDNA from the previous step. Samples should have been quantified in the previous Qubit Quantification step.

1. Use the spreadsheet template excel file to easily estimate the volume per reaction of each reagent. Template excel can be downloaded here: https://github.com/biomobot/SC2
2. Remove the pooled and AMPure XP bead cleaned up samples from the previous step and let it thaw. Keep the cDNA on ice throughout this protocol.
3. Remove NEB Ultra II End-Prep Enzyme (End-Prep Enzyme) from -20 °C, gently flick the tube several times, spin briefly and place directly on ice.
4. Remove NEB Ultra II End-Prep Reaction Buffer (End-Prep Buffer) from -20 °C and thaw at room temperature. Once thawed, vortex the reaction buffer for 10 seconds and place on ice.
   a. Make sure that the contents of each tube are clear of any precipitate and are thoroughly mixed before setting up the reaction.

5. While the reagents thaw, turn on the thermal cycler to 20 °C for 10 min, 25 °C for 5 min 65°C for 10 min, 65°C for 5 min and hold at 4 °C.
6. Set up a PCR Plate that can be used in the thermal cycler and label the wells for each pooled PCR product that will be end-prepped.

**Note:** Alternatively, PCR tubes or strips can be used to end-prep the samples if there are fewer number of end-prep reactions.

7. In a microcentrifuge tube, make a 96 sample Master Mix by combining the following and mix by pipetting up and down 5 times to mix well. The resulting Master Mix will now be referred to as the End-Prep Master Mix.

| Reagent | Volume |
|---|---|
| **Ultra II End-prep Reaction Buffer** | 196 µL |
| **Ultra II End-Prep Enzyme Mix** | 84 µL |
| **Total** | 280 µL |

**Note:** If there are less than 96 samples to be end-preped, use the excel sheet mentioned in step 1 to estimate the volume for the number of samples you have. You can also perform a calculation to determine the amount of master mix necessary for the number or samples.

8. Add the following reagents to the reaction plate/tube in the same order as shown in the table below.
   a. After adding each reagent to the reaction, mix by pipetting up and down 3 times. Be sure to switch tips between each sequential addition.
   b. When performing 96 end-prep reactions use a P10 multichannel pipette to add 2.5 µL of the End-Prep Master Mix to the reaction.

| Reagent | Volume |
|---|---|
| **ddH$_2$O** | 12-x µL |
| **cDNA** | x µL (65 ng cDNA) |
| **End-Prep Master Mix** | 2.5 µL |
| **Total** | **15 µL** |

*The x is the volume from the pooled PCR sample that yields 65 ng.

9. Seal the plate with an adhesive or a heat seal. Make sure that each well is well sealed to prevent evaporation.
   a. If using individual PCR tubes, close the lids tightly and briefly spin.
10. Spin the plate briefly at 1000 RPM for 20 seconds.
11. Place the PCR plate/PCR tubes in a thermal cycler and start the program (to 20 °C for 10 min, 25 °C for 5 min 65°C for 10 min, 65°C for 5 min and hold at 4 °C).
    a. The program will run for ~25 minutes.
12. Label the leftover cDNA plates as and place in -20 °C.
13. Once the End Prep reaction is finished the end prep reaction can be stored in 4 °C for a week or -20 °C for long term storage.

Notify the following team members that you have completed this protocol:

- Library DNA Barcoding

### Working Protocol S8: Sample Barcoding.

#### Introduction

The barcoding reaction is done using the 96 Native Barcoding Expansion 96 (EXP-NBD196). The barcoding ends contain overhangs which can be ligated to the adapter (AMII). Up to 96 samples can be multiplexed and sequenced at once on a single flow cell. Samples are first barcoded, then pooled and cleaned up using SPRI beads.

#### Materials & Reagents

**Note:** All reagents & materials were estimated for 96 samples.

##### Reagents

- Native Barcoding Expansion 96 (EXP-NBD196)
- NEBNext Ultra II Ligation Master Mix
- NEBNext Ultra II Ligation Enhancer
- AMPure XP Beads
- Short Fragment Buffer (SFB)

##### Materials

- 96-well PCR plate
- Magnetic Rack
- Multichannel Pippetes
- Plate Seal
- Thermal cycler
- Trough

##### *Barcoding Master Mix*

**Note:** Barcoding should be done on a clean bench to avoid any chance of contamination. Therefore, decontaminate your bench with RNAse decontamination solution and then 70% ethanol.

1. Thaw the native barcodes at room temperature, enough for one barcode per sample.
2. Once thawed vortex the Native Barcoding Expansion 96 (EXP-NBD196) plate and spin down at 2,000 rpm for 30 seconds.
3. In a 2 mL microcentrifuge tube, make a 96 sample Master Mix by combining the following and mix by pipetting up and down 5 times to mix well. The resulting Master Mix will now be referred to as the Barcode Master Mix.

| Reagent | Volume |
|---|---|
| **ddH2O** | 601.92 µL |
| **Ultra II Ligation Master Mix** | 264 µL |
| **Ultra II Ligation Enhancer** | 31.68 uL |
| **Total** | 1689.6 µL |

4. Mix the master mix well by pipetting up and down.
5. Add the following reagents to the reaction plate/tube in the same order as shown in the table:

a. After adding each reagent to the reaction, mix by pipetting up and down 3 times. Be sure to switch tips between each sequential addition.
b. When performing 96 end-prep reactions use a P10 multichannel pipette to add 2.5 μL of the End-Prep Master Mix to the reaction.

| Reagent | Volume |
|---|---|
| **End-prepped DNA** | 1.5 μL |
| **Native barcode** | 2.5 μL |
| **Barcode Master Mix** | 16 μL |
| **Total** | **20 μL** |

6. Mix well by pipetting up and down.
7. Seal the plate with an adhesive or a heat seal. Make sure that each well is well sealed to prevent evaporation.
    a. If using individual PCR tubes, close the lids tightly and briefly spin.
8. Spin the plate briefly at 1000 RPM for 20 seconds.
9. Place the PCR plate in a thermal cycler and start the program (to 20 °C for 20 min, 25 °C for 5 min, 65°C for 10 min, 65°C for 5 min and hold at 4 °C).
10. Label the leftover End Prep plate and place in -20 °C.
11. Once the barcoding reaction is finished remove the seal and pool all the reactions into 2 mL microcentrifuge tube.
12. Mix the contents well and spin down briefly.
13. Split the reaction equally into 2 microcentrifuge tubes.

**Note:** Reactions must be split into 2 tubes due to the volume. If barcoding less samples use 1 tube and follow the same directions. Elution volume for 1 tube should be 30 uL.

14. Resuspend the AMPure XP beads by vortexing for 30 seconds.
    a. The solution should be a homogenous brown color.
15. Check the volume of each microcentrifugre tube that contains the split reaction using a pipette.
16. Mix the beads by pipetting up and down 3 times and add 0.5x the volume of AMPure XP beads to the each of the tubes.
17. Mix the beads with sample by pipetting up and down 5 times.
18. Close the tube tightly and flick the sample several times.
19. Incubate the tube at room temperature for 10 minutes on a Hula mixer (rotator mixer).
20. Pulse centrifuge the tubes and place on the magnetic rack for 5 minutes.
    a. Be careful when opening the tube lids as the sample could travel up the wall and prevent pelleting.
21. Check whether the solution became clear. If not, continue to keep on the magnet.
22. Keep the tube on the magnet and carefully aspirate and discard the supernatant without disturbing the beads.

**CAUTION:** Be very careful not to touch the pelleted beads when removing the supernatant. Removing any of the beads will drastically decrease yield.

23. Remove the tube from the magnet and place it on the benchtop.

24. Add 500 μL of Short Fragment Buffer (SFB) to each of the tubes, pipette up and down and flick the beads to resuspend.
25. Pulse centrifuge the tubes and place on the magnetic stand for 4 minutes.
26. Check whether the solution became clear. If not, continue to keep the plate on the magnet.
27. Keep the tubes on the magnet and carefully pipette off and discard the supernatant without disturbing the beads. This ends the first SFB wash.

**CAUTION:** Be very careful not to touch the pelleted beads when removing the supernatant. Removing any of the beads will drastically decrease yield.

28. Repeat the steps for a second SFB wash.
29. Pulse centrifuge the tubes and place on the magnet.
30. Use a P10 single channel pipette to remove any residual SFB you observe.
    a. Be very careful not to remove any beads.
31. Keep the tube on the magnet and add 100 μL of freshly made 80% ethanol.
32. Remove the ethanol, spin down and use a P10 single channel pipette to remove any residual ethanol.
33. Allow to dry for ~30 sec-1 min but do not over dry the pellet.
34. Remove the tubes from the magnetic stand and onto the benchtop.
35. Flick the Elution Buffer (EB) several times and pulse centrifuge.
36. Using a P20 pipette set to 15 μL, add 15 μL of the Elution Buffer (EB) to the tube to elute the DNA and mix by pipetting up and down 5 times.
37. Incubate at room temperature for 5 minutes.
38. Flick the tubes several times and pulse centrifuge.
39. Place the tubes on magnetic stand for 5 minutes.
40. Using a P10 pipette, pool both of the elutions into a newly labeled 1.5 mL microcentrifuge tube. You should now have 30 μL of pooled barcoded sample.
41. Label, date and place the tubes immediately on ice for the next step or store at a 4 °C.

Notify the following team members that you have completed this protocol:

- Adapter Ligation

### <u>Working Protocol S9</u>: Adapter Ligation.

#### Introduction

Adapter ligation results in the addition of sequencing adapters at each end of the fragment. The barcoded pooled sample should contain a cohesive end which is used as a hook to ligate sequencing adapters. Both the template and complement strands carry the motor protein which means both strands are able to translocate the nanopore.

#### Materials & Reagents

**Note:** All reagents & materials are estimated for 96 samples.

##### Reagents

- Short Fragment Buffer (SFB) (400 µL)
- Elution Buffer (EB) (15 µL)
- Adapter Mix II (AMII) (5 µL)
- Quick Ligation Reaction Buffer 5x (10 µL)
- Quick T4 DNA Ligase (5 µL)
- AMPure XP Beads (30 µL)

##### Materials

- Magnetic Stand

#### Procedure

**Note:** If a 96 Native Barcoding Kit was used, 1 pooled sample (from the previous barcoding step), will be quantified, adapter ligated and sequenced.

**Note:** Adapter ligation should be done on a clean bench to avoid the chance of contamination. Therefore, decontaminate your bench with RNAse decontamination solution and then 70% ethanol.

##### *Qubit Quantification*

Before Adapter ligation reaction, refer to the <u>Qubit Quantification Protocol</u> and quantify each sample tube using 1 µL of sample. If using 96 barcoding kit, there will be 1 sample to quantify. Remove Qubit DNA HS reagents from 4 °C refrigerator and let them reach room temperature.

##### *Adapter Ligation Reaction*

1. Thaw Elution Buffer (EB) and NEBNext Quick Ligation reaction Buffer (5x) at room temperature. Mix by vortexing, pulse centrifuge using a benchtop centrifuge and place on ice.
   a. Make sure that the contents of each tube are clear of any precipitate and are thoroughly mixed before setting up the reaction
2. Gently flick the Quick T4 Ligase and the Adapter Mix II (AMII) several times, spin down and immediately place on ice.
3. Take the tube with 30 µL of pooled barcoded sample from previous step and add the following reagents in the same order as shown in the table below. Mix each reagent prior to adding by pipetting up and down 3 times.

   a. After adding each reagent to the reaction, mix by pipetting up and down 3 times. Be sure to switch tips between each sequential addition.

| Reagent | Volume |
|---|---|
| **Pooled Barcoded Sample** | 30 uL |
| **Adapter Mix II (AMII)** | 5 uL |
| **NEBNext Quick Ligation Reaction Buffer (5X)** | 10 uL |
| **Quick T4 DNA Ligase** | 5 uL |
| **Total** | **50 uL** |

4. After adding Quick T4 DNA Ligase, mix the contents of the tube by flicking the tube. Briefly centrifuge using a tabletop centrifuge.
5. Close the tubes tightly and incubate the reaction at room temperature for 25 minutes.
   a. While the reaction is incubating you can aliquot AMPure XP beads (next section).

##### *AMPure Bead Clean Up*

**Note:** Aliquot 500 µL from the stock AMPure XP into a labeled and dated 1.5 mL microcentrifuge tube and use this for AMPure clean up instead of the large stock bottle. Vortex the commercial stock bottle for 30 seconds prior to aliquoting into the 1.5 mL microcentrifuge tube. Repeat aliquoting as needed.

1. Resuspend the 1.5 mL microcentrifuge tube containing AMPure XP beads by vortexing for 30 seconds.
   a. The solution should be a homogenous brown color.
2. Mix the beads by pipetting up and down 3 times and add 30 µL of AMPure XP beads to the tube containing the pooled barcoded samples from the previous step.
3. Mix the beads with sample by pipetting up and down 5 times.
4. Close the tube tightly and flick the sample several times.
5. Incubate the tube at room temperature for 10 minutes on a Hula mixer (rotator mixer).
6. Pulse centrifuge the tubes and place on the magnetic rack for 5 minutes.
   a. Be careful when opening the tube lids as the sample could travel up the wall and prevent pelleting.
7. Check whether the solution became clear. If not, continue to keep on the magnet.
8. Keep the tube on the magnet and carefully aspirate and discard the supernatant without disturbing the beads.

**CAUTION:** Be very careful not to touch the pelleted beads when removing the supernatant. Removing any of the beads will drastically decrease yield.

9. Remove the tube from the magnet and place it on the benchtop.
10. Add 200 µL of Short Fragment Buffer (SFB) to each of the tubes and flick the beads to resuspend.
11. Pulse centrifuge the tubes and place on the magnetic stand for 4 minutes.
12. Check whether the solution became clear. If not, continue to keep the plate on the magnet.
13. Keep the tubes on the magnet and carefully pipette off and discard the supernatant without disturbing the beads. This ends the first SFB wash.

**CAUTION:** Be very careful not to touch the pelleted beads when removing the supernatant. Removing any of the beads will drastically decrease yield.

14. Repeat 10-13 for a second SFB wash.
15. Pulse centrifuge the tubes and place on the magnet.
16. Use a P10 single channel pipette to remove any residual SFB you observe.
    a. Be very careful not to remove any beads.
17. Remove the tubes from the magnetic stand and onto the benchtop.
18. Flick the Elution Buffer (EB) several times and pulse centrifuge.
19. Using a P20 pipette set to 15 µL, add 15 µL of the Elution Buffer (EB) to the tube to elute the DNA and mix by pipetting up and down 5 times.
20. Incubate at room temperature for 5 minutes.
21. Flick the tube several times and pulse centrifuge.
22. Place the tubes on magnetic stand for 5 minutes.
23. Using a P10 pipette, carefully transfer the final library to a new labeled 1.5 mL microcentrifuge tube.
24. Label, date and place the tubes immediately on ice for the next step or store at a 4 °C.

Notify the following team members that you have completed this protocol:

- Quantify, Load and run MinIONs

### Working Protocol S10: MinION Loading and Running.

#### Introduction

Flow cells are shipped with storage buffer to maintain product integrity. The priming step flushes out the storage buffer and replaces it with a mix of Flush Buffer (FB) and Flush Tether (FLT). The DNA library is then loaded into the flow cell. The sequencing run is monitored in real time using MinION software and Rampart via fast basecalling.

#### Materials & Reagents

**Note:** You may refer to a video on how to load the flow cell: https://youtu.be/Pt-iaemrM88

##### Reagents

- Loading Beads (LB) - 25.5 µL
- Sequencing Buffer (SQB) - 37.5 µL
- Flush Buffer (FB) - 1 mL
- Flush Tether (FLT) - 30 µL
- Qubit Reagents (Refer Qubit Protocol)

##### Materials

- SpotON Flow Cell
- MinION

#### Procedure

##### *Qubit Quantification*

**Note:** Prior to priming and loading the flow cell, refer to the Qubit Quantification Protocol and quantify the sample from previous step using 1 µL of sample. If using 96 barcoding kit, there will be 1 sample to quantify. Remove Qubit DNA HS reagents from 4 °C refrigerator and let them reach room temperature. You will load 20 ng onto the flow cell in 12 µL volume.

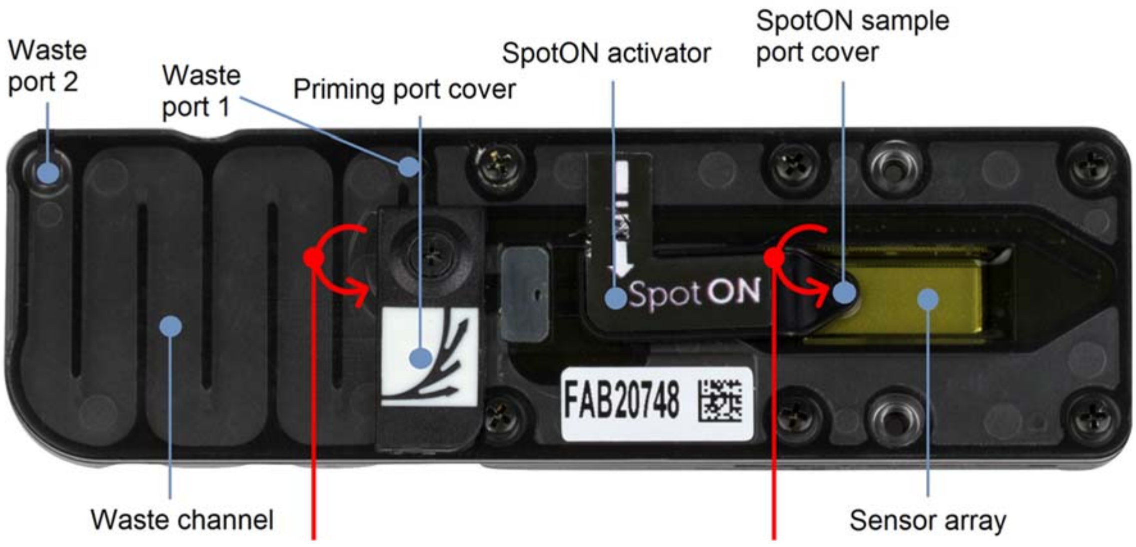


**Figure 1:** MinION Flow Cell

#### *Prepare Priming Mix*

**Caution:** The priming step flushes out the storage buffer and replaces it with a mix of Flush Buffer (FB) and Flush Tether (FLT). The DNA library is then loaded into the flow cell. Throughout the process it is **essential** that the sensor array remains submerged in buffer at all times. **If an air bubble passes over any channels, those pores will be permanently damaged.**

1. Thaw the Sequencing Buffer (SQB), Loading Beads (LB), Flush Tether (FLT) and one new tube of Flush Buffer (FB) taken from -20 °C at room temperature before placing the tubes on ice as soon as thawing is complete.
2. Mix the Sequencing Buffer (SQB), Flush Buffer (FB) and Flush Tether (FLT) tubes by vortexing, spin down, then return to ice.
3. Open the MinION Sequencer lid and slide the flow cell under the clip.
   a. Press down firmly to on the flow cell to ensure correct thermal and electrical contacts.
4. Transfer 30 μL of Flush Tether (FLT) directly into a new tube of Flush Buffer (FB). Label the FB tube lid with an asterisk, to indicate FLT has been added.
5. Mix the contents of the tube by flicking the tube with your finger and then spin down briefly in a microfuge.
   a. The resulting buffer is now referred to as the **Priming Mix** (FB + FLT). You will use this to flush out the storage buffer from the flow cell.

#### *Priming the Flow Cell*

1. Open the priming port by sliding the cover clockwise so that the port is visible as shown below.

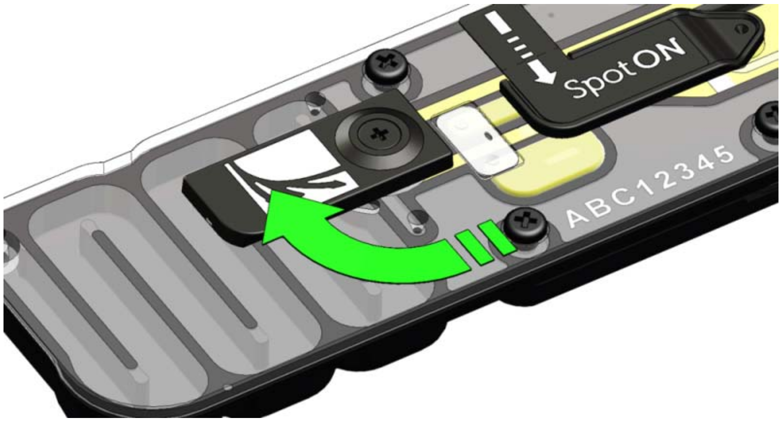


**Figure 2:** Opening the priming port on the flow cell.

**Caution:**

When drawing up solution:

- make sure there is no air plug at the bottom of the pipette tip

When ejecting the solution:

- do not fully expel the liquid from the tip
- leave a small volume in the tip end so that no air can follow the solution into the priming port

2. After opening the priming port, draw back a small volume to remove any bubbles (~20 µl):
    a. Set a P1000 pipette to 200 µl
    b. Insert the tip into the priming port
    c. Turn the wheel until the dial shows 220-230 µl, or until you can see a small volume of buffer entering the pipette tip.
3. With the priming port open, use a P1000 pipette, to slowly add ~800 µl of priming mix.
    a. Leave a tiny bit of priming mix in the pipette tip to avoid introducing air bubbles.
4. Close the priming port and wait 5 minutes. While waiting, prepare the pre-sequencing mix found below.

#### *Prepare Pre-Sequencing Mix*

**Note:** The Loading Beads (LB) tube contains a suspension of beads. These beads settle very quickly. It is vital that they are mixed immediately before use by pipetting up and down. Dilute the DNA library if necessary, using Elution Buffer (EB) to 20 ng in 12 uL.

1. In a new tube, prepare the Pre-Sequencing Mix for loading as follows:

| Reagent | Volume |
|---|---|
| **Sequencing Buffer (SQB)** | 37.5 µl |
| **Loading Beads (LB), mixed immediately before use** | 25.5 µl |
| **DNA library** | 12 µl |
| **Total** | **75 µl** |

2. Mix the contents of the tube by flicking the tube and spinning down briefly.
3. Place on ice until steps for priming of the flow cell have been completed.

#### *Cont. Priming the Flow Cell*

1. Open the priming port by sliding the cover clockwise so that the port is visible as shown in figure 2.
2. Gently lift the SpotON sample port cover to make the SpotON sample port accessible. You will still need to use the priming port, so keep this open.
3. Using a P1000 pipette, slowly load 200 µl of the Priming Mix (FB mixed with FLT) into the priming port.

**Caution:** Do NOT eject to the last stop of the pipette, as this would introduce air bubbles. Pipette slowly, to avoid the buffer bubbling out of the sample port. Carry out this step immediately before loading the library.

4. Resuspend the Pre-Sequencing Mix by gently pipetting it up and down using a P200 pipette and tip, to get a homogenous mixture.
    a. Pipette up and down carefully to avoid creating air bubbles. Gentle resuspension also helps prevent accidental shearing of the DNA.
5. Mix the prepared library gently by pipetting up and down just prior to loading.
6. Add 75 µl of sample to the flow cell via the SpotON sample port in a dropwise fashion. Ensure each drop flows into the port before adding the next.

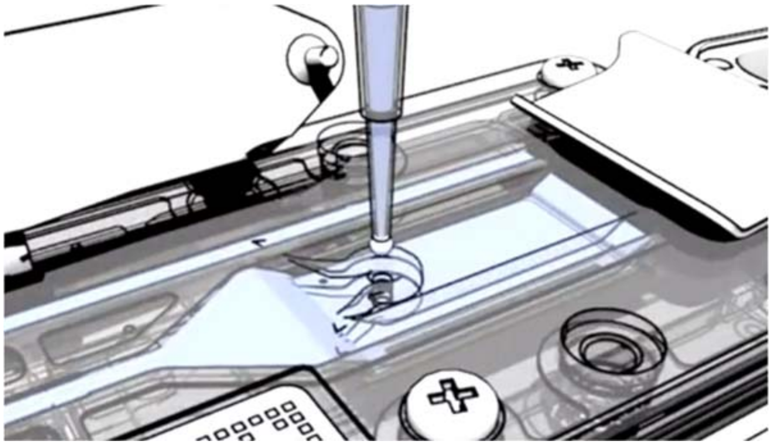

**Figure 3:** Loading the library onto the SpotOn flow cell.

7. Close the SpotON port with the cover ensuring the bung enters into the port.
8. Close the priming port.
9. Close the MinION lid

#### *Starting a Sequencing Run*

1. Plug in the MinION into the computer's USB 3.0 port also known as the SS port.
2. Double click the MinION software to open it.
3. In the Connection Manager window click **This Device.**
4. Click on **Jump to run** in the lower left-hand side.
5. Flow cell type will automatically come up as FLO-MIN106. If not, select **FLO-MIN106**.
6. Click on **New Experiment** at the bottom left of the window. A new window will appear.

**Experiment**

- Name of experiment starting with the date. Example: **YYMMDD_sc2_run**

**Kit**

- Select **SQK-LSK109** as the kit
- For Barcoding Expansion Packs do **NOT** select a barcoding kit

***Basecalling***

- Select Basecall model to **Fast Basecalling.**
- Barcoding **off**.

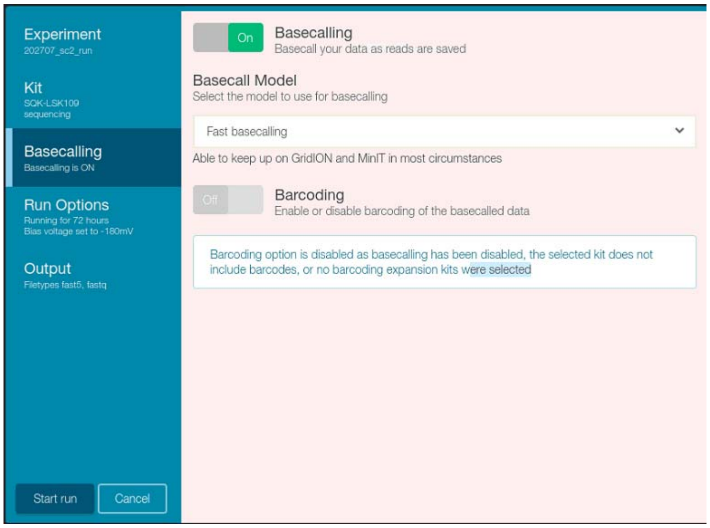

**Run Options**

- Leave as default.

**Output**

- Select the output location to be **/home/gagnonlab/sc2_seq_data**
- Select the FAST5 and FASTQ Reads per File to be **1000.** Click on **Start Run.**

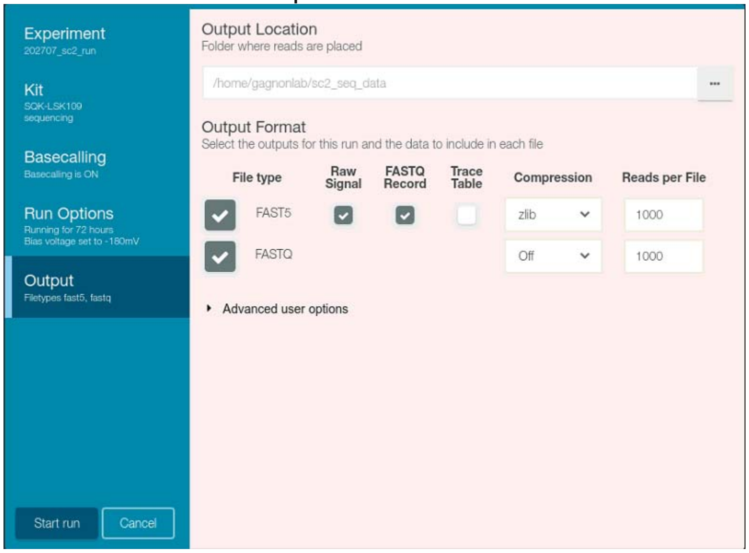

### **Working Protocol S11**: Flow Cell Wash and Store.

#### Introduction

The Wash Kit allows sequential runs of multiple sequencing libraries on the same flow cell. It works by washing out the first library, and refreshing the system ready for a subsequent library to be loaded. This procedure provides the opportunity to utilize the same flow cell a number of times, maximizing the available run time, particularly for cases where less data per library is required. Following the wash step, Storage Buffer can be introduced into the flow cell, allowing storage of the flow cell before subsequent library additions.

#### Materials & Reagents

##### Reagents

- Wash Solution A
- Wash Solution B
- Storage Buffer

##### Materials

- Ice
- Flow Cell

##### *Preparation to Wash and Store*

| Contents | Tube Volume | Per Wash/Store Use |
|---|---|---|
| **Wash solution A** | 140 µL | 20 µL |
| **Wash Solution B** | 1400 µL | 380 µL |
| **Storage Buffer** | 1800 µL | 500 µL |

1. Remove **Wash Solution A** from -20 °C, gently flick the tube, spin briefly and place directly on ice.
2. Remove **Wash Solution B** from -20 °C and thaw at room temperature. Once thawed, vortex, briefly spin and place on ice.
3. Remove **Storage Buffer** from -20 °C and thaw at room temperature. Once thawed, mix by pipetting up and down, briefly spin and place on ice.
4. Stop sequencing run by opening MinKnow software and selecting STOP sequencing. Do NOT stop the Basecalling and leave MinKnow software open.
   a. The software will continue to basecall the raw sequenced data.
5. Once the sequencing run has been stopped, leave the flow cell in the device, unplug the MinION and carefully place the MinION device on your benchtop.

##### *Wash and Store*

1. In a clean microcentrifuge tube, prepare the following **Wash Mix**:

| Component | Volume |
|---|---|
| **Wash solution A** | 20 µL |
| **Wash Solution B** | 380 µL |

2. Mix well by pipetting, spin briefly and place on ice.
   a. Do not vortex the tube.

3. Ensure that the priming port cover and SpotON sample port cover are closed, in the positions indicated in the figure below.

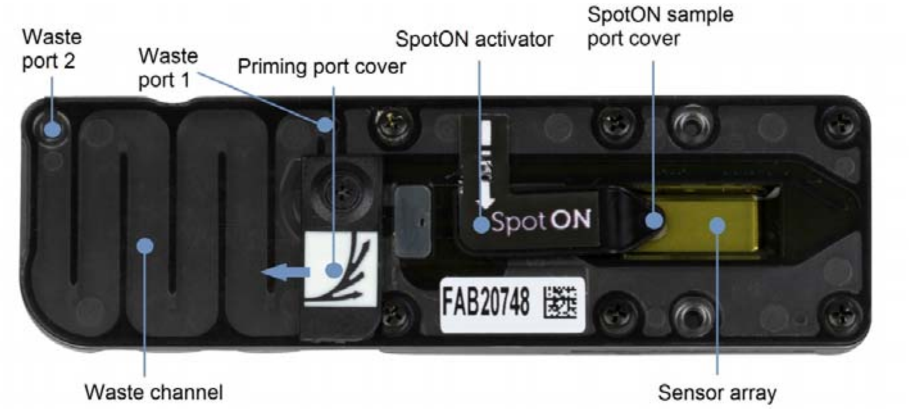


4. Using a P1000 set to 1000 µL, remove all fluid from the waste channel through **Waste port 1**. As both the priming port and SpotON sample port are closed.

**Important:** It is vital that the flow cell priming port and SpotON sample port are closed to prevent air from being drawn across the sensor array area, which would lead to a significant loss of sequencing channels.

5. Rotate the flow cell priming port cover clockwise so that the priming port is visible.
6. Using a P1000 set to 200 µl draw back a small volume to remove any air.
    a. Insert the tip into the priming port.
    b. Turn the wheel until the dial shows 230 µl. Small volume of liquid may enter the pipette tip.
    c. Dispense the liquid and leave the priming port open.
7. Using a P1000 set to 400 µl, load 400 µl of the prepared **Wash Mix** into the flow cell via the priming port, avoiding the introduction of air.

**Important:** Be careful not to introduce air into the flow cell when adding the Wash Mix. Do not load the full 400 µl, leave a small amount of Wash mix in the pipette tip to avoid introducing air bubbles.

8. Close the priming port, close the MinION Lid and wait for 25 minutes.
9. Once 25 minutes has passed, ensure that the priming port cover and SpotON sample port cover are closed.
10. Using a P1000, remove all fluid from the waste channel through **Waste port 1**. As both the priming port and SpotON sample port are closed.
11. Rotate the flow cell priming port cover clockwise so that the priming port is visible.
12. Slowly add 500 µl of Storage Buffer (S) through the **priming port** of the flow cell avoiding the introduction of air bubbles.
13. Close the **priming port**.
14. Using a P1000, remove all fluid from the waste channel through **Waste port 1**. As both the priming port and SpotON sample port are closed
15. Carefully remove the flow cell from the MinION device and place it in the clear plastic tray. Keep the flow cell horizontal the entire time.
16. Place the clear plastic tray in the original packaging and seal the packaging. Keep the flow cell horizontal.

17. Label the original packaging with the Experiment Name, Today's Date, Number of Pores, and Hours Sequencing. All of this can be found on MinKNOW software under "Experiments" and "System Messages."

18. The flow cell can now be stored at 4 °C for next use.

### Working Protocol S12: Sequencing Data Analysis.

##### *Program Installation*

Guppy Basecaller and Barcoder: https://community.nanoporetech.com/downloads

pycoQC: https://github.com/a-slide/pycoQC

ARTIC Bioinformatics environment setup: https://artic.network/ncov-2019/ncov2019-it-setup.html

ARTIC Bioinformatics Pipeline: https://artic.network/ncov-2019/ncov2019-bioinformatics-sop.html

##### *Folder structure*

Once the sequencing is completed you will see a folder in the directory saved with the latest sequencing run.

Make a folder structure that is easy to navigate to. Example shown below:

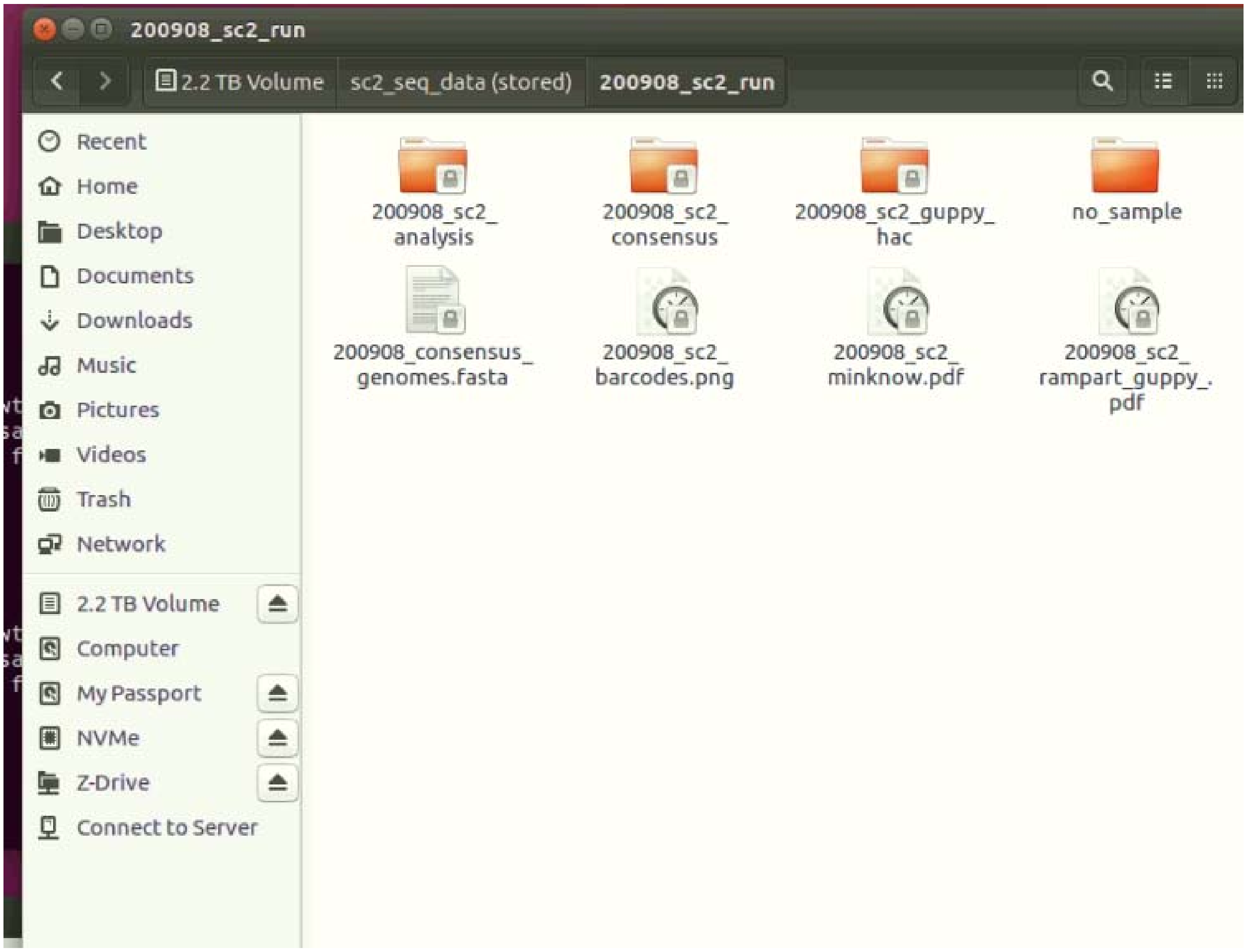


This is an example of good folder structure for a single batch.

##### *Guppy GPU Basecalling*

Once the folders structure is set up the data will be basecalled from the fast5 raw signal files to fastq files using Guppy_Basecaller.

Open the terminal using the keys **CTRL+SHIFT+T**.
In the terminal enter the following command making sure that the input and output directories are correct.

Example:

```
guppy_basecaller --input_path
/home/gagnonlab/sc2_seq_data/YYMMDD_sc2_run/fast5 --save_path
/home/gagnonlab/sc2_seq_data/YYMMDD_sc2_run/YYMMDD_sc2_guppy_hac --config
dna_r9.4.1_450bps_hac.cfg --device cuda:0
```

> It is possible to have several different directories of fast5 files. Ex: fast5_skip fast5_pass, fast5_fail. In that case all the fast5 files should be combined into a single directory and guppy_basecaller should be ran on all the fast5 files generated from the sequencing run. By default reads with a mean q-score value greater than 7 are kept which corresponds to roughly 85% basecall accuracy.

If successful, the following should display in the terminal with the 0-100% indicating progress of basecalling.

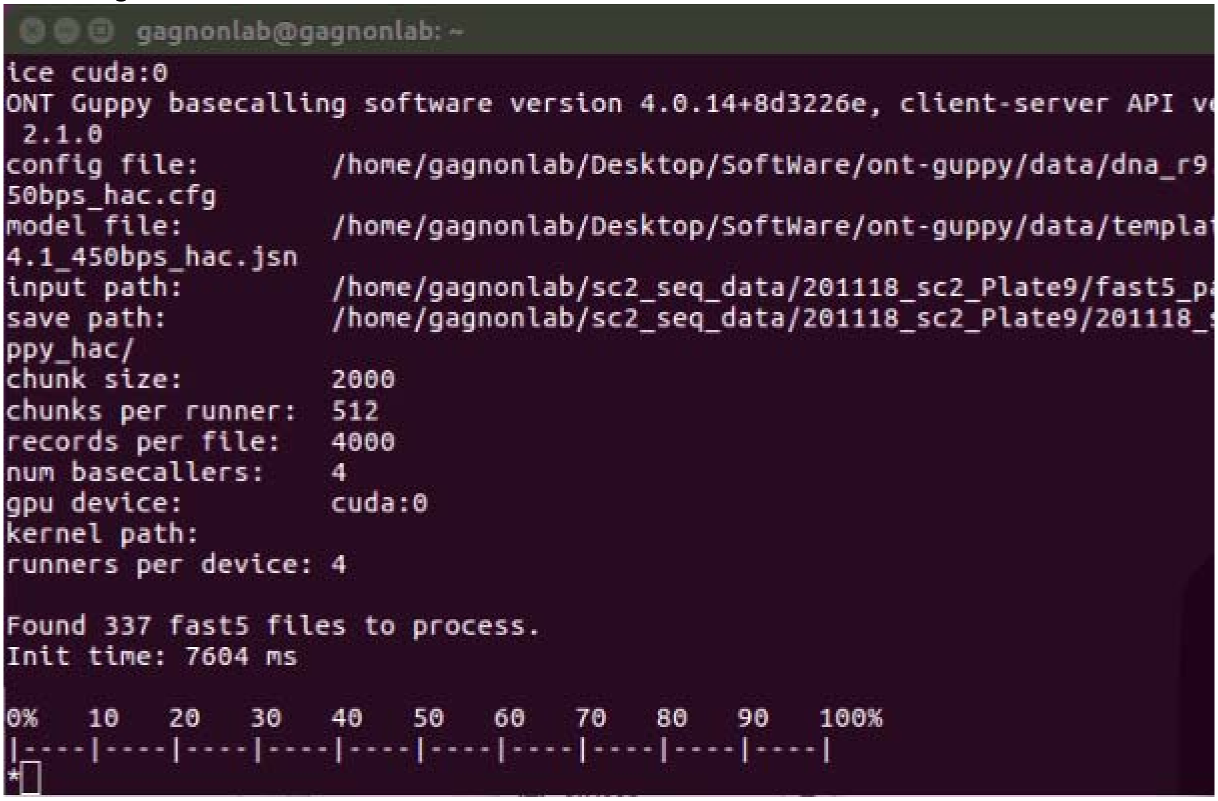


Once guppy_basecaller has basecalled all of the fast5 files, you should see them in the output directory.

##### *QC the Data (optional)*

Sequencing metrics of the run could be generated by using pycoQC.

```
pycoQC -v -f
/home/gagnonlab/sc2_seq_data/201118_sc2_Plate9/201118_sc2_guppy_hac/sequencin
g_summary.txt -o 201118_pycoQC.html
```

The above command will create the file run_1.html with multiple plots and summary statistics.

#### *Guppy Demultiplexing*

To demultiplex your basecalled fastq files use the guppy_barcoder.

-i Your input will be the guppy basecalled directory of the fastq files
-s Your save directory will be the directory you have made for your demultiplexed files

Enter the following command. It is important that –require barcodes both ends is selected.

```
guppy_barcoder --require_barcodes_both_ends -i
/home/gagnonlab/sc2_seq_data/YYMMDD_sc2_run/YYMMDD_sc2_guppy_hac -s
/home/gagnonlab/sc2_seq_data/YYMMDD_sc2_run/YYMMDD_sc2_demultiplex --
arrangements_files barcode_arrs_nb96.cfg
```

*if you are using a different barcoding kit than the 96 native barcoding kit (EXP-NBD196) the –arrangements_files will change.

For the 1-12 and 13-24 barcoding kit use the following command:

```
guppy_barcoder --require_barcodes_both_ends -i
/home/gagnonlab/sc2_seq_data/YYMMDD_sc2_run/YYMMDD_sc2_guppy_hac -s
/home/gagnonlab/sc2_seq_data/YYMMDD_sc2_run/YYMMDD_sc2_demultiplex --
arrangements_files barcode_arrs_nb12.cfg barcode_arrs_nb24.cfg
```

If successful, the following should display in the terminal with the 0-100% indicating progress of basecalling.

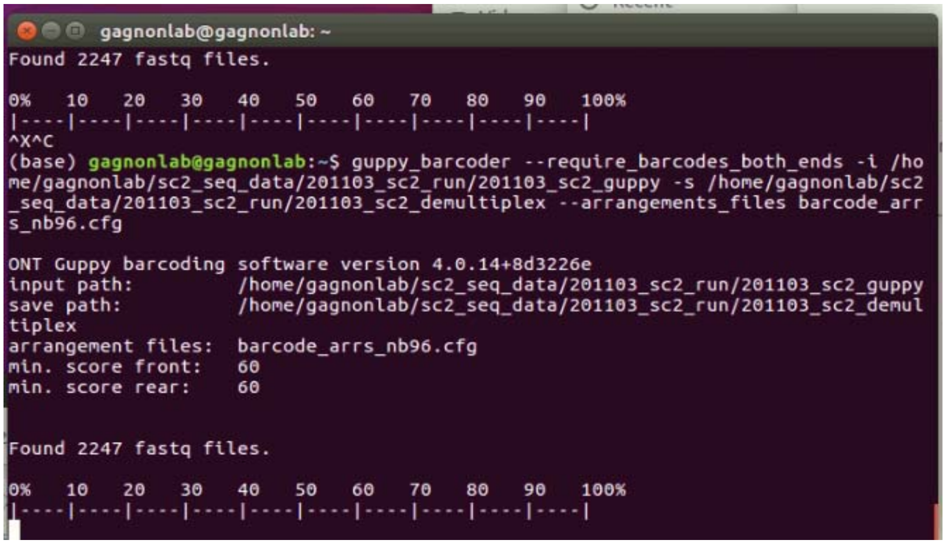

***Rampart***

To determine the coverage and generate RAMPART PDF you can enter the following command in a new terminal. This can be done on the Guppy demultiplexed data or concurrently with MinKNOW.

Guppy demultiplexed Rampart basecalledPath: the output directory of guppy_demultiplex.
MinKNOW concurrently Rampart basecalledPath: the Fastq_pass directory of your MinKNOW sequencing run.

To begin activate the artic-ncov2019 conda environment:

```
conda activate artic-ncov2019
```

Then enter the following command to start Rampart:

```
rampart --clearAnnotated --protocol /home/gagnonlab/artic-ncov2019/rampart --basecalledPath
/home/gagnonlab/sc2_seq_data/YYMMDD_sc2_run/YYMMDD_sc2_demultiplex/
```

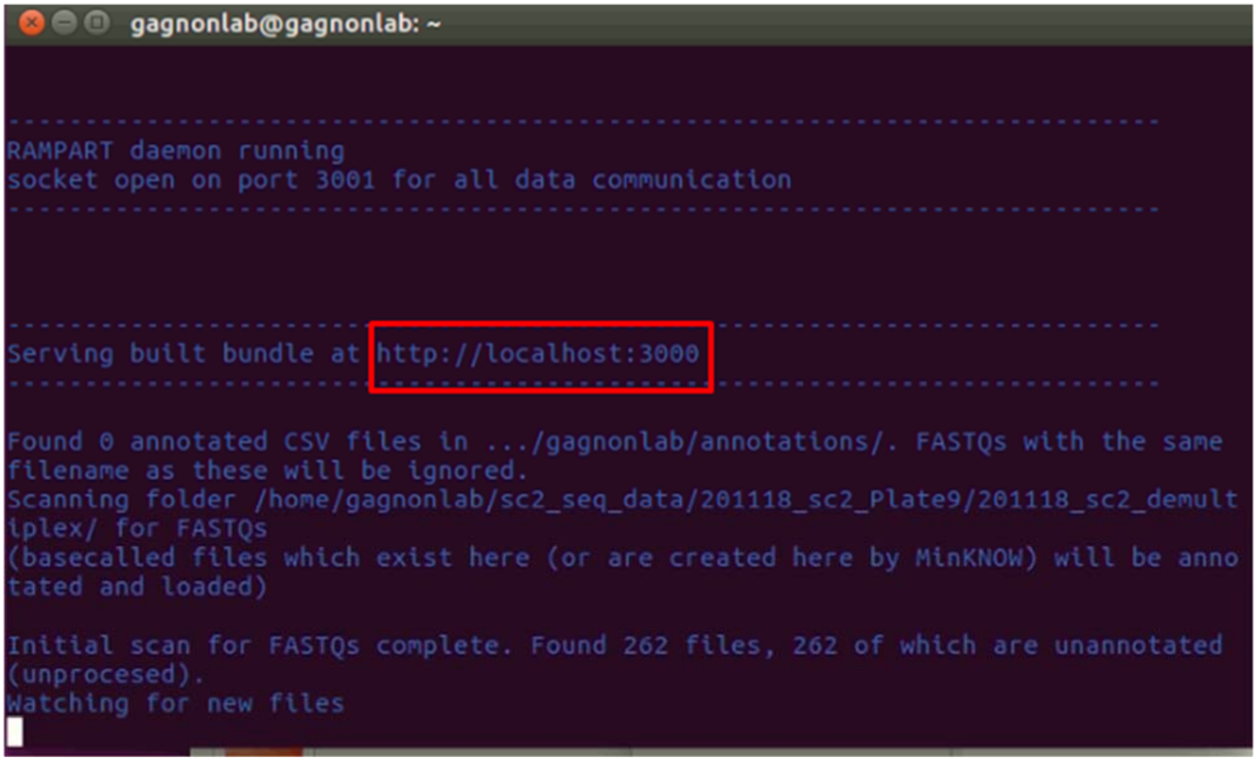


Open a browser at localhost:3000 where data processing can be visualized.

#### *ARTIC Guppyplex*

To filter reads by length reads, navigate into a directory called YYMMDD_sc2_filtered. This is where all your filtered fastq will be outputted.

Open the terminal if not already opened. Enter the following command:

```
conda activate artic-ncov2019
```

Navigate to the YYMMDD_sc2_filtered directory by using the `cd` and `ls` commands. Once in this directory you will filter the samples one at a time for the barcodes that have reached the desired >20x percent coverage.

* The 20x % coverage for each barcode can be determine by using Rampart.

Enter the following command to filter the reads between 400-700 bp:

```
artic guppyplex --min-length 400 --max-length 700 --directory
/home/gagnonlab/sc2_seq_data/YYMMDD_sc2_run/YYMMDD_sc2_demultiplex/barcodeXX
--prefix XX_NBXX_YYMMDD_sc2_filtered
```

*Make sure you have the barcode that matches the sample for each filtering that is performed.

***ARTIC Medaka Pipeline***

To run the Medaka pipeline run the following command with the available threads in your system:

```
artic minion --medaka --normalise 0 --threads 12 --scheme-directory
/home/gagnonlab/artic-ncov2019/primer_schemes --read-file
XX_NBXX.fastq nCoV-2019/V3 XXX_NBXX
```

A genome consensus fasta file should be generated at the end of the pipeline run. Any poition that is not covered by at least 20 reads from either read group are mased as ambiguous bases (Ns).

***Concatenate all FASTA files into One File:***

```
cat *.consensus.fasta > YYMMDD_consensus_genomes.fasta
```

### Working Protocol S13: Visualizing Data in Nextclade and Nextstrain.

### Sequence preparation and cleaning

Before inserting sequences into a NextStrain pipeline, it is most efficient to remove those that will not pass Nextstrain quality control. Open a web-browser and navigate to https://clades.nextstrain.org/. Nextclade is a simple site to observe the mutations, gaps, and general quality of your sequences.

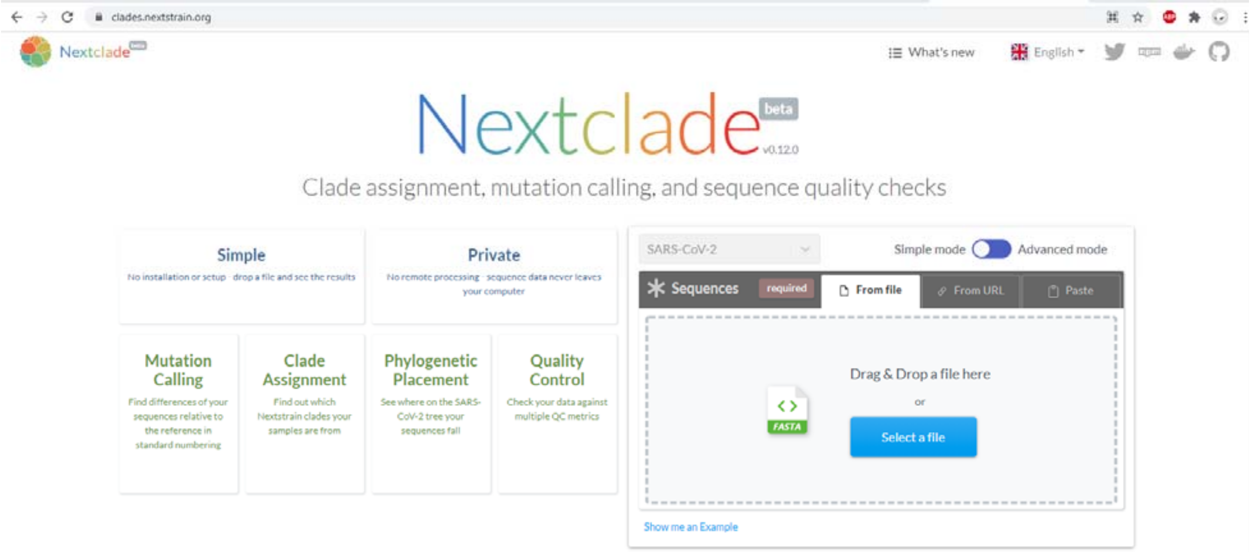


There is a box on this page labeled 'Drag & Drop a file here'. Provide this box with the FASTA file of your new sequences. This will initiate an automated procedure where each sequence will be analyzed.

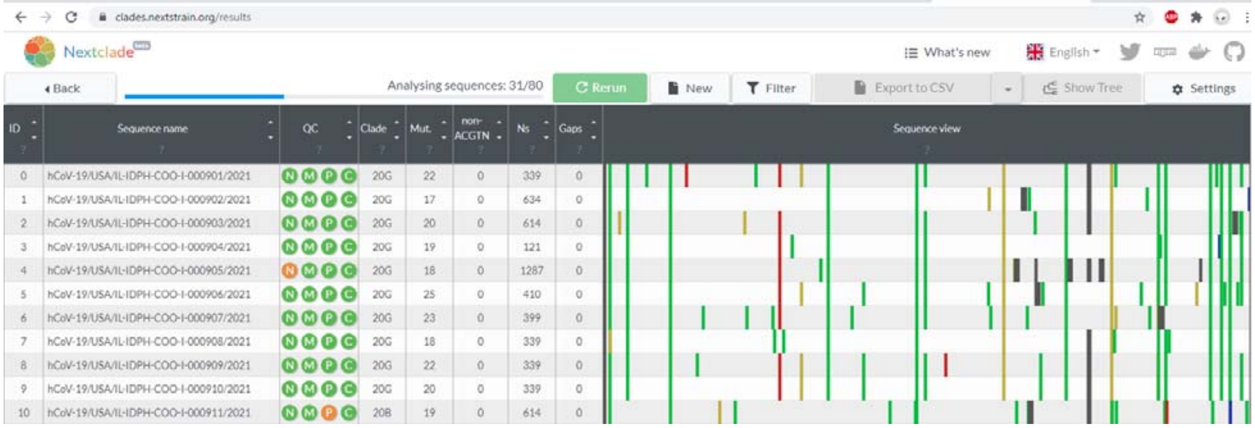


Each column provides different information about your sequence:

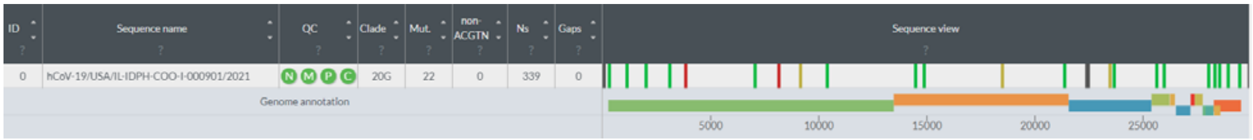

- Sequence name – name provided for sequence in FASTA file
- QC – General indications of quality of sequence. In general, green and yellow are acceptable, red is unacceptable. Hovering the mouse over each circle gives more details.
  - N circle – measurement of missing data. Based on the number of Ns in the sequence. Will be flagged (yellow) with more than 1000 Ns. Will fail (red) with more than 3000 Ns.
  - M circle – measurement of mixed sites. Will be flagged if more than 10 nucleotides have mixed states (such as R or Y), as this may be indicative of contamination.
  - P circle – measurement of private mutations. Based on the number of mutations present compared to the original Wuhan-Hu-1 sequence.
  - C circle – measurement of mutation clusters. Based on the clustering of mutations. If more than 6 mutations are within a 100 base range, the sequence will be flagged.
- Clade – What Nextstrain clade the software determines the sequence to be. This label is reliable with good quality reads, but may be unreliable with too many missing nucleotides.
- Mut. – Number of mutations in the sequence compared to original SARS-CoV-2 sequence, Wuhan-Hu-1. Hovering the mouse over a specific sequence's mutation column will generate a popup window with a list of nucleotide and amino-acid mutations.
- Non-ACGTN – Number of ambiguous, non-N nucleotides in the sequence. Hovering the mouse over a specific sequence's Non-ACGTN column will generate a list.
- Ns – Number of missing nucleotides in the sequence. Hovering the mouse over a specific sequence's Ns column will generate a list.
- Gaps – Number of intentional gaps in the sequence (-). Produced by deletions in the nucleotide code. Hovering the mouse over a specific sequence's Gaps column will generate a list.
- Sequence View – Color-coded visual representation of the mutations, gaps, and Ns in the sequence, as compared to Wuhan-Hu-1. Red, Mutation to A. Blue, Mutation to C. Yellow, mutation to G. Green, mutation to T. Dark grey, N. Light grey, gap. A visual representation of SARS-CoV-2 genes is provided below the sequences for comparison. Hovering the mouse over a specific color-coded bar will provide additional information about it.

In addition, there are several useful tools at the top of the page:

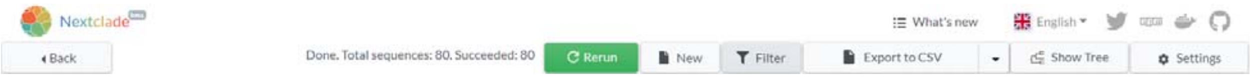


- Filter – Filter your sequence list based on various criteria, including a mutation at a specific nucleotide or amino-acid, the quality of the sequence, or the Nextstrain clade it is categorized as.
- Export to CSV – Allows the generation of a spreadsheet with all details of the analysis for later browsing.
- Show Tree – places sequences on a phylogenetic tree for comparison with other sequences from the GISAID Initiative.
- Settings – Adjust settings for each of the four quality control determinations.

Using these criteria, poor-quality sequences can be identified and removed from your FASTA file.

### Acquisition of global SARS-CoV-2 data

Sequences and metadata for global SARS-CoV-2 sequences can be acquired from a number of sources. Nextstrain assumes, by default, the use of data from the GISAID Initiative (https://www.gisaid.org/). After registering an account, a compressed file can be downloaded for the most recent metadata and FASTA sequences. At the time of this writing, that is in excess of 450,000 sequences.

### Metadata Preparation

Metadata must be prepared for each sequence that will be put through the Nextstrain pipeline. Prepare a tab-separated values (.tsv) file for your data. This is also known as a tab-delineated file. It should conform to the same column order and formatting as the metadata file provided by GISAID. This order could be changed later, but at the time of this writing, it is:

**strain** – sequence name. Just exactly match the sequence name in the FASTA file.
virus – “ncov”
gisaid_epi_isl – GISAID ID
genbank_accession – Genbank ID
**date** – date of sample collection. Must be in YYYY-MM-DD format.
**region** – continent of sample collection.
**country**- country of sample collection.
**division** - division of sample collection. A more granular location, such as a state within the United States of America.
location – most granular location of sample collection. Often a county or city.
**region_exposure** – continent of patient exposure, if known. If not known, identical to region.
**country_exposure** – country of patient exposure, if known. If not known, identical to country.
**division_exposure** – division of patient exposure, if known. If not known, identical to division.
segment – “genome”
**length** – full length of sequence. Is used to remove insufficiently long sequences.
**host** – species of sample collection patient
**age** – age of sample collection patient. If unknown, enter “?”
**sex** – sex of sample collection patient. If unknown, enter “?”
Nextstrain_clade – Nextstrain-determined clade. Typically consists of a number, signifying the year it appeared, followed by a letter.
pangolin_lineage – PANGO lineage, as determined by the Pangolin tool
GISAID_clade – Clade as determined by GISAID
**originating_lab** – lab that collected the sample from the patient
**submitting_lab** – lab that sequenced the genome
authors - submitting lab head, typically
url – URL associated with author or submitting lab
title – Title of paper associated with submitted genomes
paper_url – URL of paper, if any
**date_submitted** – Date submitted to GISAID
purpose_of_sequencing – Currently under-used. May be used in the future to indicate samples that were S-gene dropouts, for example.

Necessary items are in bold. However, even if a column is blank, a space must still be made for it, so that it can be integrated into a larger metadata file more easily.

### Installing Nextstrain

Nextstrain can be installed as detailed in the tutorial (https://nextstrain.github.io/ncov/). These instructions are for Unix-based systems. Start with a system that has Conda installed. Execute the following commands:

```
curl http://data.nextstrain.org/nextstrain.yml --compressed -o
     nextstrain.yml
conda env create -f nextstrain.yml
conda activate nextstrain
npm install --global auspice
```

This creates a build environment and a folder called `ncov` that can be used to run the Nextstrain pipeline.

### Putting sequence data into Nextstrain

By default, the FASTA file and metadata file for the sequences are located in the `data` subdirectory. If using global data from GISAID, name the files `sequences.fasta` and `metadata.tsv`, respectively. To add the lab's own data to this repository, commands can be used to append the lab's data to the end of these files. For example, if the lab's FASTA file is named `lab.fasta`, the following command could be used:

```
cat lab.fasta >> sequences.fasta
```

Lab metadata can be appended to the end of the global metadata file, using the following command, which leaves out the first line, which should the header:

```
tail -n +2 -q lab.tsv >> metadata.tsv
```

### Setting up a profile

In order for the Nextstrain pipeline to function as required, it must know what location your analysis is focusing on, and otherwise how to select a subsampling of its sequences. Example profiles are in the `nextstrain_profiles` subdirectory. It is recommended to copy one of these example profiles to the `my_profiles` subdirectory and modify it to meet your needs. Within each profile folder is several files:

config.yaml – this configuration file points to the location of further files, and tells Nextstrain how verbose to be and how many CPU cores to use.

builds.yaml – This file tells nextstrain how to subsample the sequences to build its phylogenetic tree. It is not practical to use all 400,000+ samples. A 3000-5000 subsample tree, on the other hand, can easily be built in an afternoon, depending on computer resources. Here is an example builds.yaml that collects every sample in the state of interest (Illinois), and then a small subsample from other states, other countries, and other continents.

```
builds:
  illinois:
    subsampling_scheme: illinois
    geographic_scale: division
    region: North America
    country: USA
    division: Illinois
    title: "Illinois Samples with contextual global samples"

subsampling:
  illinois:
    # Focal samples for division
    division:
      group_by: "year month"
      seq_per_group: 99999999
      exclude: "--exclude-where 'region!={region}'
     'country!={country}' 'division!={division}'"
    # Contextual samples from division's country
    country:
      group_by: "division year month"
      seq_per_group: 5
      exclude: "--exclude-where 'region!={region}'
     'country!={country}' 'division={division}'"
      priorities:
        type: "proximity"
        focus: "division"
    # Contextual samples from division's region
    region:
      group_by: "country year month"
      seq_per_group: 4
      exclude: "--exclude-where 'region!={region}'
     'country={country}'"
      priorities:
        type: "proximity"
        focus: "division"
    # Contextual samples from the rest of the world, excluding
     the current
    # division to avoid resampling.
    global:
      group_by: "country year month"
      seq_per_group: 3
      exclude: "--exclude-where 'region={region}'"
      priorities:
        type: "proximity"
        focus: "division"
```

Briefly, the build name, recorded on the second line, determines the name of the final json file output by the Nextstrain pipeline. In this example, the build name is Illinois, so the resulting file is `ncov_illinois.json`. Whatever the subsampling scheme is labeled in line three, it must be expanded upon in the subsampling section, starting on line ten. In this example, the Illinois subsampling scheme is broken down into four further subsamplings: division, country, region, and global. The division section says to sort all samples by year and month, exclude those that are outside Illinois, and then select all of the resulting selection. The country section says to sort all samples by division, year, and month, and then exclude those outside the United States, as well as those inside Illinois. From the remaining divisions of the United States, select at random, five sequences per month per division. This pattern continues in the region and global sections.

Other files in the profile can similarly be altered to customize, e.g., the colors or formatting of the final output of the Nextstrain pipeline, but the defaults can be used for now, until specific customizations are required.

### Other files to modify

Other files in the ncov directory may need to be modified to accommodate the lab's data:

defaults/lat_longs.tsv – A tab-delineated file that contains the latitude and longitude of all regions, countries, divisions, and locations. If the lab's data includes any divisions or locations that are no in the global nextstrain build yet, they can be added manually here.

defaults/clades.tsv – A tab-delineated file that determines what nucleotide or amino-acid mutations are defined as a clade in the phylogenetic tree. This file is updated regularly by Nextstrain, but if additional clades are desired, they can be added manually here.

### Running the profile

Once everything is set up, all that is left to do is run the Nextstrain pipeline. Nextstrain uses the `snakemake` command to initiate the pipeline. If a custom profile has been set up, it needs to be invoked at this state. For example, the 'illinois' profile mentioned above is used via the following command:

```
snakemake --profile ./my_profiles/illinois
```

The pipeline will begin to run. This can take anywhere from several minutes to several hours to several days, depending on computational resources and size of the subsample used. Briefly, the samples are first filtered to remove any too-short sequences. Then the remaining sequences are aligned and masked to better compare them. Next, subsampling is done using the criteria set up in the profile. After this, a tree is built, and then refined using the dates in the metadata. Amino-acid mutations are calculated based on nucleotide mutations, and clades are determined. These outputs accumulate in the results subdirectory. Particularly time-intensive steps include the MAFFT step, in which all sequences are aligned, and the tree refining step. While the former can utilize all available CPUs, and is only long due to the sheer number of sequences in the GISAID dataset, the latter is currently only optimized for one CPU at a time, and can run for quite a long time if the subsampled sequence size is too large.

### Visualizing the result

Once the Nextstrain pipeline completes, a JSON file will be created in the auspice subdirectory. This file can be viewed by opening a web browser to [https://auspice.us/](https://auspice.us/) and dragging-and-dropping the JSON file onto it. This will generate a completed nextstrain profile.

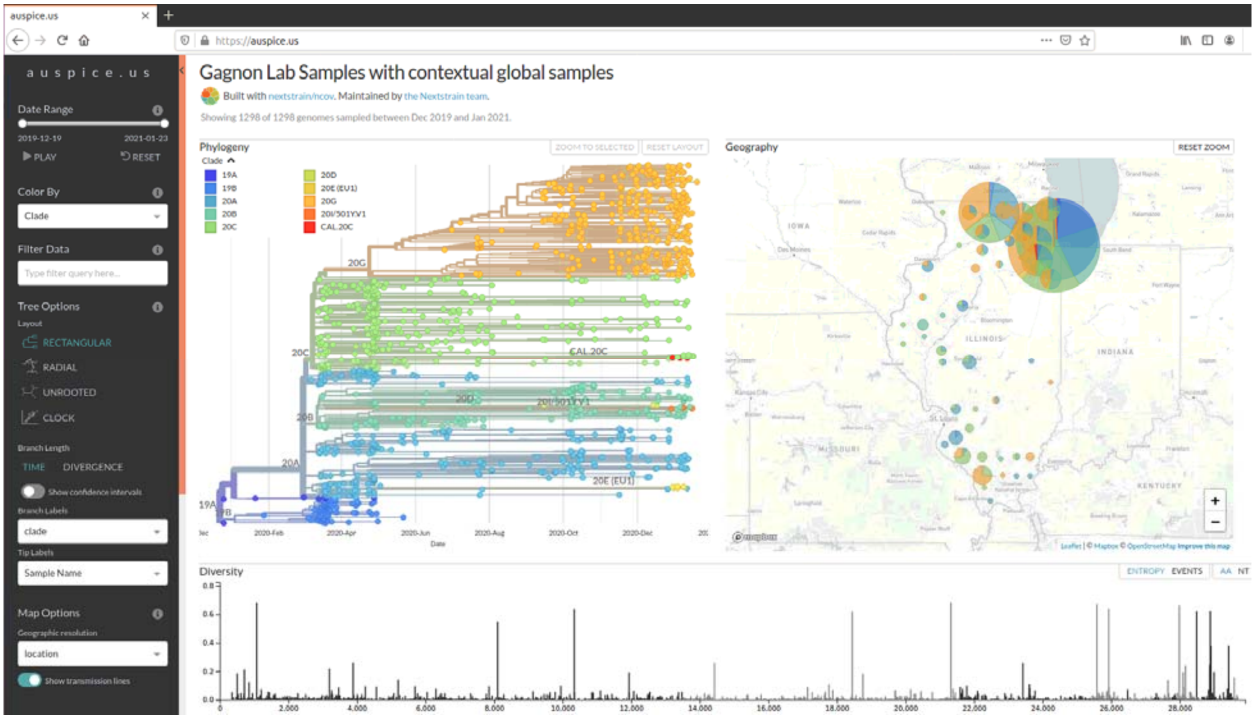


This interface allows a great number of display options and visualizations, including coloring samples based on different criteria (clade, specific mutations, location, data of collection, etc). It can also be set to display only samples from specific locations, clades, data, etc, using the filters at the bottom on the page.
